# A method to study extracellular vesicles secreted *in vitro* by cultured cells with minimum sample processing and extracellular vesicle loss

**DOI:** 10.1101/2021.06.18.447964

**Authors:** A Viveiros, V Kadam, J Monyror, LC Morales, D Pink, A M Rieger, S Sipione, E Posse de Chaves

## Abstract

Extracellular vesicles (EVs) are involved in a multitude of physiological functions and play important roles in health and disease. The study of EV secretion and EV characterization remains challenging due to the small size of these particles, a lack of universal EV markers, and sample loss or technical artifacts that are often associated with EV separation techniques. We developed a method for in-cell EV labeling with fluorescent lipids (DiI), followed by DiI-labelled EV characterization in the conditioned medium by imaging flow cytometry (IFC). Direct IFC analysis of EVs in the conditioned medium, after removal of apoptotic bodies and cellular debris, significantly reduces sample processing and loss compared to established methods for EV separation, resulting in improved detection of quantitative changes in EV secretion and subpopulations compared to protocols that rely on EV separation by ultracentrifugation. In conclusion, our optimized protocol for EV labeling and analysis reduces EV sample processing and loss, and is well suited for cell biology studies that focus on modulation of EV secretion by cells in culture.

## Introduction

Extracellular vesicles (EVs) are cell-derived particles delimited by a membrane. They are shed by most cells and play pivotal roles in intercellular communication and signaling both in health and disease conditions (Mathews and Levy, 2019; Murao et al., 2020; Raposo and Stahl, 2019; Srivastava et al., 2021). Further to this, EVs can carry cancer biomarkers and therefore have diagnostic and prognostic value (Hu et al., 2020; Kim et al., 2020). There is also a growing appreciation of the utility of EVs as therapeutics (Born et al., 2020; Nam et al., 2020; Sarko and McKinney, 2017; Zocchi et al., 2020), gene editing tools (Kostyushev et al., 2020), drug carriers (Mehryab et al., 2020) and in other clinical applications (Andjus et al., 2020; Yan et al., 2020). Owing to this multitude of physiological functions and therapeutic applications, the interest in EVs has grown exponentially in the past few years and recent technological advances have considerably expanded the tools available for EV studies. Nevertheless, the small and heterogenous size (30-1000 nm) of EVs and the absence of universal EV markers are still challenges that significantly hamper studies on the biology of EVs, with many aspects related to biogenesis, secretion, and biology still remaining unclear (Margolis and Sadovsky, 2019; Stahl and Raposo, 2019).

Recent surveys indicated that the largest proportion of studies on EVs are based on the analysis of EVs secreted in the cell culture conditioned medium (Gardiner et al., 2016; Royo et al., 2020). The most common method of EV separation is ultracentrifugation alone or combined with other techniques. Ultrafiltration-size exclusion chromatography (SEC) has been gaining popularity in the past 5 years (Royo et al., 2020). A variety of methods are used to characterize EVs including Western blotting, electron microscopy, single particle tracking (mostly nanoparticle tracking analysis (NTA), and flow cytometry (Gardiner et al., 2016; Royo et al., 2020).

In order to gain deeper insights into the mechanisms that underlie EV secretion and other aspects of EV biology, improved methods for EV separation and analysis are required. Characterization of EV subpopulations that might express different markers or carry different cargos is also of paramount importance, as biochemical analysis of bulk EV preparations may not adequately reveal quantitative and qualitative differences in EV subpopulations. Significant strides in this direction have been made through the development of methods for single particle analysis by high-resolution flow cytometry (van der Vlist et al., 2012) and imaging flow cytometry (IFC). A comprehensive analysis for applying this technology to the study of EVs is available (Erdbrugger et al., 2014; Gorgens et al., 2019; Lannigan and Erdbruegger, 2017). Characterization of EVs by IFC requires the use of fluorescent markers because EVs display very low level of scatter and most cannot be detected in brightfield images because the resolution of the BF camera is not sufficient (Lannigan and Erdbruegger, 2017). EV labeling for these studies is most often performed after separation from conditioned medium or biological fluids, an approach that presents with several challenges, including elimination of fluorescently-labeled non-EV particles and/or free-dye(Lai et al., 2015; Morales-Kastresana et al., 2017; Puzar Dominkus et al., 2018). Moreover, although several dyes are already in use for EV labeling, there is a recognized need for improved dyes and labeling methods (Russell et al., 2019).

Here, we aimed to design a simplified and reliable method to study EV secretion and regulation in cell models. In this study, *i)* we present a method to label EVs that consists of labeling parental cells with a lipophilic cationic indocarbocyanine dye prior to EV collection and analysis by IFC; *ii)* we demonstrate that the cleared conditioned medium (depleted of cell debris and apoptotic bodies) represents the preferred sample for analysis, with minimum processing and loss compared to EV preparations that rely on ultracentrifugation or size exclusion chromatography; and *iii)* we provide evidence that, at least in some cases, EV analysis performed in the cleared conditioned medium allows detection of changes in EV secretion that cannot be distinguished if the samples undergo ultracentrifugation.

## Materials and Methods

### Materials

Opti-MEM^®^ reduced-serum medium, Dulbecco’s Modified Eagle Medium (DMEM), penicillin/streptomycin, fetal bovine serum (FBS), N-2 supplement, B27 supplement and Geneticin^®^ were from Gibco. Protease inhibitor cocktail tablets were from Roche. Clarity™ Western ECL Substrate and nitrocellulose membranes were purchased from Bio-Rad. Bicinchoninic acid (BCA) protein assay kit was from Thermo Fisher Scientific. Vybrant™ DiI was from Invitrogen. All horseradish peroxidase-conjugated secondary antibodies were from Amersham Biosciences. qEV columns (SP1) were from Izon (Christchurch, New Zealand). Immobilon PVDF membranes, Amicon Ultra-15 100K MWCO, Amicon Ultra-4 10K MWCO and Amicon Ultra-0.5 10K MWCO filters, NP-40 (IGEPAL^®^ CA-630) and Bafilomycin A1 were purchased from Millipore Sigma. Glutathione Sepharose 4B beads were purchased from GE Healthcare. Penicillin/Streptomycin were purchased from Hy-Clone™, GE Healthcare. Pre-lubricated pipette tips (Maxymum Recovery™, Axygen^®^,1-200μL and 100-1000μL) and microcentrifuge tubes were purchased from Corning Life Sciences.

### Cell culture

Mouse neuroblastoma Neuro2a (N2a) cells were cultured in DMEM/Opti-MEM I (1:1) supplemented with 5% heat-inactivated fetal bovine serum (FBS) and 1% penicillin/ streptomycin. Cells were maintained at 37°C with 5% CO_2_ and split every 3-4 days. Medium was changed every second day.

### Fluorescent labeling of cells for EV detection

N2a cells were labelled with the lipophilic membrane stain DiI (1,1’-dioctadecyl-3,3,3’,3’-tetramethylindocarbocyanine perchlorate; λEx/λEm = 549/565 nm) according to the manufacturer’s instructions (Invitrogen™, Thermo Fisher Scientific, USA). Briefly, cells were detached with 0.25% trypsin-EDTA, resuspended in serum-free medium (Opti-MEM:DMEM (1:1), 1% penicillin/streptomycin, 2mM L-glutamine and 0.11g/l sodium pyruvate) to a concentration of 1×10^6^ cells/ml. Five μl of the dye solution (1mM) was added to each 1ml of cell suspension and incubated for 20 min at 37°C in the dark and then centrifuged at 300 x *g* for 5 min at r.t. The stained cell pellet obtained was further subjected to three rounds of centrifugation in growth media to remove unbound dye. Cells were plated at 1.8×10^6^ cells/10cm diameter plate in DMEM/Opti-MEM I (1:1) supplemented with 5% FBS, 1% penicillin/streptomycin, 2mM L-glutamine and 0.11g/l sodium pyruvate, and maintained in culture for 18 h before starting EV collection. When using dishes of other sizes the seeding densities were determined such that the confluence of the cells at the time of harvesting was ~ 80%. For EV collection, the medium was replaced with medium containing N-2 supplement, but no FBS (serum- and phenol red-free medium), and the conditioned medium was harvested after 24 h.

### Collection of EVs in cell-conditioned media

After cell treatment, medium was replaced with serum-free media supplemented with N-2 supplement (OptiMEM:DMEM 1:1, 1% penicillin/streptomycin, 2mM L-Glutamine and 0.11g/L Sodium Pyruvate with 1x N-2),and allowed to be conditioned by cells for 24 h. The conditioned medium was collected in polypropylene tubes and centrifuged at 2,000 x g for 10 min at 4°C in an Eppendorf^®^ Centrifuge 5810 R, using an A-4-81 swinging bucket rotor, to pellet any remaining cells, apoptotic bodies, and cell debris, resulting in a cleared conditioned medium. The cells were harvested in 500 μl of PBS and lysed by sonication. Protein concentration in cell lysates was measured with a Pierce™ BCA Protein Assay and or a Pierce™ Enhanced BCA Protein Assay (Thermo Fisher Scientific, USA) according to manufacturer’s instructions. Before further processing, cleared conditioned media from different samples were adjusted based on cellular protein content to ensure that all cleared conditioned media derived from the same amount of cellular proteins. EV samples were always kept on ice. They were mixed by tube inversion or by gently pipetting up and down and analyzed immediately. All culture media and diluents were prepared fresh and filtered through a 0.1 μm filter.

### Fluorometry

DiI fluorescence was measured in cell lysates and in the cleared conditioned medium using SpectraMax^®^ i3x multi-mode microplate reader (Molecular Devices, USA). λEx/λEm for DiI is 540/580nm. The bandwidth of all excitation and emission wavelengths was set to 15nm. Fluorescence was measured in 96-well black-bottom plates by well-scan reading. Fifty microliters of the sample were loaded in each well in triplicates.

### Separation of EVs by sequential ultracentrifugation

EV separation by sequential ultracentrifugation (UC) was performed in accordance with previously described protocols (Lobb et al., 2015), with some modifications. Briefly, the cleared conditioned medium was centrifuged in a Beckman Coulter Optima™ MAX-XP Ultracentrifuge at 100,000 x *g* at 4°C for 90 min, using a MLA-55 fixed-angle rotor (k-factor: 53). The supernatant (Sup_100k_) was set aside at 4°C and the pellet was washed with 1ml of phosphate buffered saline (PBS) and centrifuged again at 100,000 x *g* at 4°C for 90 min, in a MLA-130 fixed-angle rotor (k-factor: 8.7). The resulting pellet (Pellet_100k_) was resuspended in 50-100 μl PBS. EVs were kept at 4°C and analyzed immediately following resuspension.

### Separation of EVs by ultrafiltration combined with size exclusion chromatography

Cleared conditioned media were concentrated using Amicon^®^ Ultra-15 Centrifugal Filters (10,000 MWCO or 100,000 MWCO) (Millipore Sigma, USA). The clear conditioned medium was loaded onto the filter and concentrated by centrifugation at 2,608 x *g* at 4°C to ≤300 μl. The concentrate was collected, and the filter membrane was washed with 100 μl serum-free medium, which was added to the concentrate. Where necessary, sample volume was adjusted to 550 μl with serum-free medium. A 50μL aliquot was used for DiI measurements by spectrofluorometry and the remaining 500μL were used for size exclusion chromatography (SEC). qEV original Size (10ml) Exclusion Columns (iZON Science^®^) were used for the separation of EVs as indicated by the manufacturer. SEC columns were stored at 4°C in PBS containing 0.05% sodium azide and used according to the manufacturer’s instructions. After column equilibration at r.t., the elution time of 10ml of PBS was recorded to ensure optimal column packing and performance. Twenty-two fractions of 0.5ml were collected using PBS as eluent. The presence of EVs in the various fractions was determined by measuring DiI fluorescence (λEx/λEm = 540/580 nm) in each fraction. Protein content of each fraction was determined by measuring the absorbance at 280nm using Nanodrop™ 2000c (Thermo Fisher Scientific, USA). The EV-rich, protein-low fractions (generally between fractions 5 and 11 included) referred to as “EV fractions” were pooled and concentrated using Amicon^®^ Ultra-2mL filters (10,000 MWCO) that were blocked with Tween-80 as previously described. EVs were maintained at 4°C and analyzed immediately.

### *In vitro* labeling of EVs

Labeling of EVs in vitro was performed by adding fluorescently-conjugated antibodies and Annexin V directly to the cleared conditioned medium or to the Pellet_100K_ resuspended in PBS. For CD9 detection we used Alexa Fluor^®^ 647-conjugated anti-CD9 antibody (BioLegend, Cat#124810) at a final concentration of 2.5μg/ml and we run in parallel samples with IgG isotype control (BioLegend, Cat#400526). All antibodies were centrifuged for 10 min at 14,000 x *g* before use. Pacific Blue™ Annexin V (Invitrogen™ A35122) was used to detect surface phosphatidylserine. Briefly, samples from the same preparation of cleared conditioned medium and Pellet_100K_ were diluted 1:1 with 2X Annexin V binding buffer (50 μl of each for a total volume of 100 μl). Next, we added 5μL of Pacific Blue™ Annexin V (1:20 dilution, concentration proprietary) and 5 μl of a 1:10 dilution of anti-CD9 in PBS. Samples were incubated at r.t. for 15 min, in the dark, prior to direct analysis by IFC. Controls including only the Annexin V label, or CD9 antibody or DiI label were used to create a compensation matrix as described in the next section. Titrations of CD9 antibody and Annexin V were performed to determine the appropriate concentrations (Supplementary Fig. 1 and 2).

### Analysis of EVs by Image Flow Cytometry (IFC)

All samples were analyzed on an ImageStreamX MkII instrument (ISX; Amnis/Luminex) equipped with 4 lasers (120.00 mW 405 nm, 200.00 mW 488 nm, 150.00 mW 642 nm, and 70.00 mW 785 nm). All lasers were set to maximum powers. All data were acquired using a 60X magnification objective, with numerical aperture of 0.9. The 60X magnification generates the lowest pixel resolution (0.3μm^2^/pixel) and will also set the core stream width to 7 μm (Gorgens et al., 2019; Lannigan and Erdbruegger, 2017). All samples were collected in the following channels: Ch03 for DiI (560-595 nm), Ch07 for Annexin V (435-505 nm), Ch11 for CD9 (642-745 nm). Ch01 (435-480nm filter) was used for bright-field imaging and Ch06 (745–780 nm filter) for SSC detection. Where indicated, the 405 laser (Ch07) was also used for SSC detection. Standard, unfiltered BioSure sheath fluid (D-PBS, pH 7.4) was used for all measurements. For each sample, acquisition was set up to capture all events that displayed lower SSC than the SpeedBeads (Amnis^®^ SpeedBead^®^ Kit for ImageStream^®^, Luminex Corp.). Unstained EVs secreted by unlabeled cells were used to assure that no fluorescence signal was detected in any channel. Similarly, the buffer only and the unconditioned N-2-supplemented medium were also run through IFC to determine the background signal. In all cases minimal to no signal events were detected. In experiments in which EVs subtypes were analyzed, single stained controls were always run in parallel to be able to establish a compensation matrix. In addition, controls with single and double antibodies or antibodies/AnnexinV in buffer were included. All samples were subjected to detergent lysis with NP-40 (0.5%) for 30 min at r.t., as described previously (Gorgens et al., 2019; Inglis et al., 2015), Data analysis was performed using Amnis IDEAS software (version 6.2). A compensation matrix was applied to address potential spectral spillover using single stained controls and the IDEAS compensation wizard. We gated on Intensity_Ch06 (SSC) in order to remove any remaining SpeedBeads from the analysis. To ensure we were analyzing only 1 EV in each image, we created a mask to identify DiI intensity (Intensity(M03,Ch03_DiI, 50-4095)). Using this mask, we developed a spot count feature (Spot Count_Intensity (M03,Ch03_DiI, 50-4095)_4) and gated on images that had no more than 1 DiI spot. The resulting population represented EVs.

A compensation matrix for spectral spillover was calculated using the single stained controls and the IDEAS_6.2 compensation wizard. The matrix was applied to all other samples, including the buffer controls, to remove spectral crosstalk between fluorochrome channels. These compensated files are then further analyzed using a variety of data analysis tools available in the IDEAS and FCS Express software (DeNovo Software). The log of the Intensity feature within the combined mask (MC) of all channels is used to plot both fluorescence and scatter parameters. The Intensity feature is the sum of all pixel values within the mask minus the background. IDEAS allows all feature values to be exported as .fcs files which can be analyzed by other flow cytometry software programs.

### Nanoparticle Tracking Analysis (NTA)

Measurement of particle size and concentration was performed by NTA, using a NanoSight LM10 system equipped with a 405nm laser (NanoSight, Amesbury, United Kingdom). EV samples were diluted with 0.1μm-filtered PBS to achieve a concentration of 20-100 particles/frame and injected into the sample chamber with sterile syringes. Five aliquots from each sample were measured, each run for 1 min. The precise temperature during sample acquisition was recorded manually to accurately determine particle concentration. All samples were captured with the same camera level and detection threshold (camera level set at 14 and detection level threshold set at 7). Instrument settings were checked prior to data collection using NIST traceable 200 nm-polystyrene beads (Thermo Scientific, 3000 series) diluted in 10mM potassium chloride. Comparison of particle concentration across different samples used dilution and volume corrected values.

### Transmission electron microscopy

Fifty microliters aliquots of the EV pellet (Pellet_100K_, freshly separated by UC and re-suspended in PBS) and SEC fractions were fixed with an equal volume of 2x Karnovsky fixative (0.2M Na cacodylate, 4% paraformaldehyde and 4% glutaraldehyde)(Ted Pella Inc.), mixed gently and incubated on ice for 30 min. Five to ten *μ*l of this mix were transferred onto carbon-formvar-coated grids (Ted Pella Inc.) following glow discharge. After 5-10 min, the grids were washed gently and transferred on top of 2.5% uranyl acetate (EMS Inc.) for 10 min. The grids were washed gently again and left to dry on a Kimwipe^®^ in the dark. Imaging was performed using a Jeol JEM-2100 microscope

### Immunoblotting

Cells were harvested in RIPA buffer (140mM NaCl, 20mM Tris, 1% SDS, 0.1% NP40, 0.5% Sodium deoxycholate, pH 7.4) supplemented with protease inhibitor cocktail and sonicated. Protein content was determined by the BCA assay (Pierce) according to manufacturer’s instructions. Absorbance at 562nm was measured using a SpectraMax^®^ i3x Multi-Mode Microplate Reader (Molecular Devices, USA). Proteins were resolved by SDS-PAGE in 16% polyacrylamide gels and transferred to PVDF membranes overnight at 4°C. Membranes were washed three times with TBS + 0.1% Tween 20 (TTBS) and blocked for 1 h in TTBS containing 5% non-fat milk. Membranes were probed overnight at 4°C with the following primary antibodies: anti-ALIX (AIP1) (1:250; BD Biosciences, 611621), anti-CD9 (1:1000; Abcam, ab92726), anti-LC3 (1:500; Novus Biologicals, NB100-2220) and anti-actin (1:2000; Cell Signaling Technology, 4957S). All primary antibodies were diluted in TTBS containing 5% bovine serum albumin (BSA), with the exception of the anti-LC3 and anti-actin antibodies, which were diluted in blocking buffer. The next day, membranes were washed twice with TBS, twice with TTBS and twice again with TBS followed by 1 h incubation with the appropriate secondary antibodies (1:2000 in blocking buffer) at r.t. with gentle agitation. Membranes were washed again twice for 5 min each with TBS, TTBS and TBS respectively. Immunoreactivity was detected with Clarity™ Western ECL Substrate (Bio-Rad) and visualized with a Li-COR C-DiGit western blot scanner. Band densitometry was performed using Image Studio (LI-COR Biosciences).

### Statistical analysis

Statistical analyses were performed in GraphPad PRISM (version 9.1.2). Comparisons between two samples were performed using paired or unpaired Student’s *t*-test. For multiple comparisons, one-way ANOVA was applied. Multiple comparisons post hoc test was performed as indicated in the figure legends.

## Results

### Cell and EV labeling

All the experiments were performed using N2a cells, which are a model extensively used to study neuronal differentiation, axonal growth and signaling pathways.

To minimize variability due to sample handling, we labeled EVs prior to their secretion by incubating parental cells with the lipophilic cationic indocarbocyanine dye DiI (Fig. 1). Cell analysis by imaging flow cytometry (IFC) demonstrates that the dye is incorporated throughout the cells and labels the plasma membrane as well as intracellular membranes, up to at least 46 h from the initial labeling (Fig. 2A). The fluorescence of stained cells was ~ 2 orders of magnitude above background autofluorescence (Fig. 2B and C). Over 95% of cells were labeled with DiI up to 46 h post-staining and throughout the timeframe of EVs collection into the conditioned medium (Fig. 2C). The overall median cell fluorescence had decreased at the latest timepoint measured (46h) compared to cells freshly stained or 22h post-staining (Fig. 2D). However, we could not detect any change of the median EV fluorescence in the timeframe of EV collection (Fig. 2E).

**Fig. 1.**
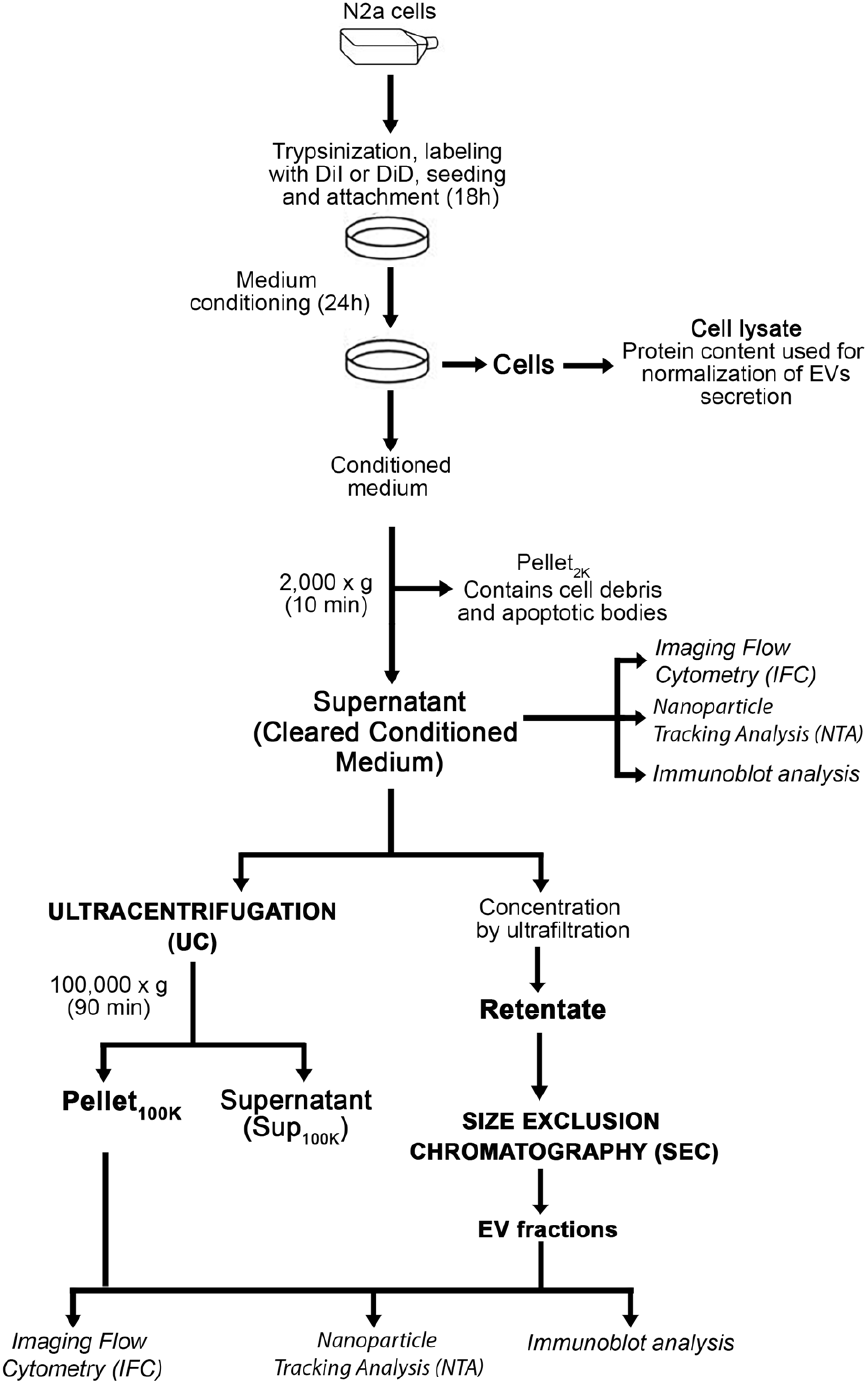
Outline of procedures for EV labeling, collection and separation. Techniques used to analyze EV samples are indicated in italic.

**Fig. 2.**
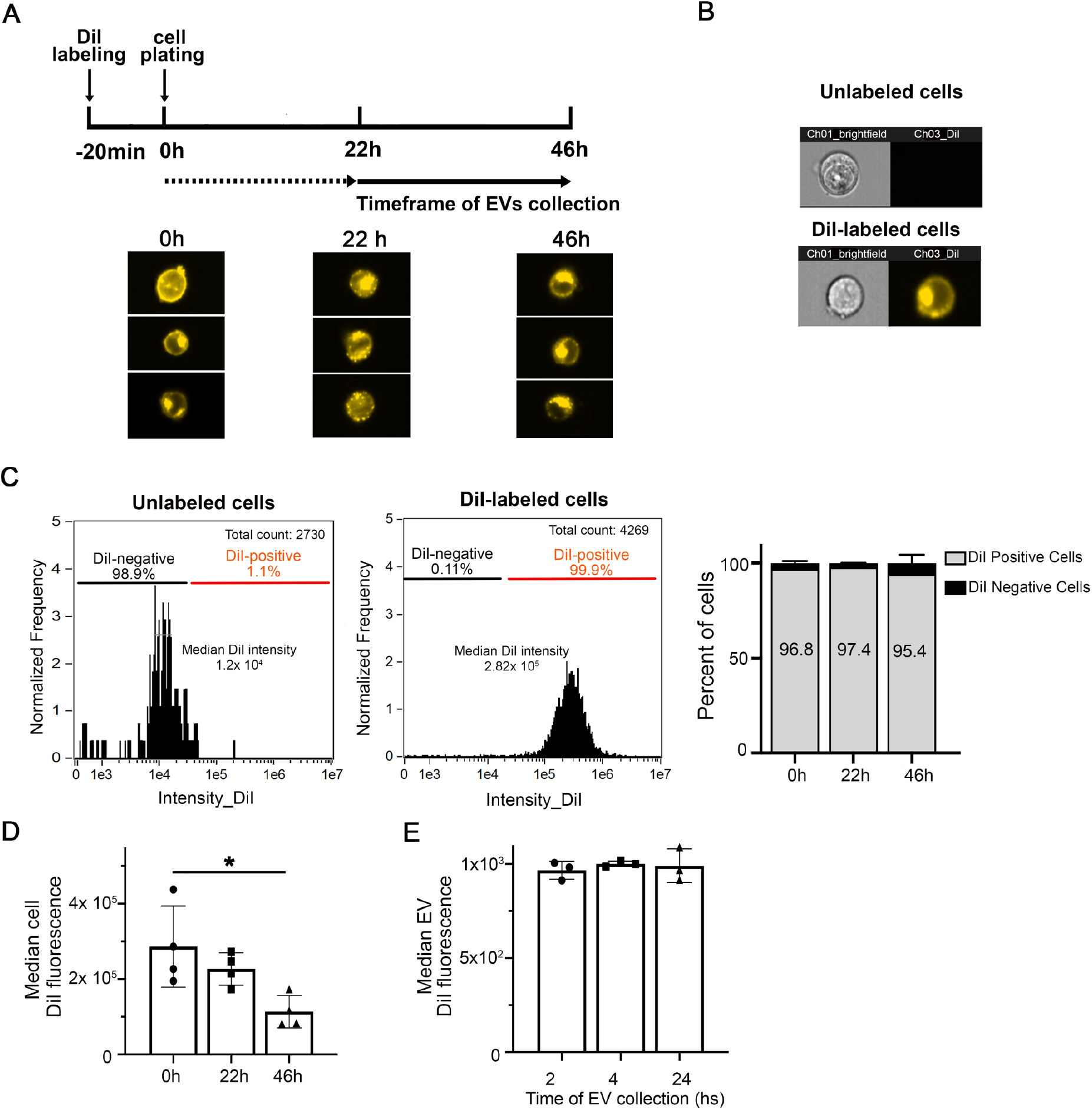
DiI effectively labels cells in culture. (**A**) Time-course of fluorescent cell analysis and EV collection after N2a cell labeling with DiI. Representative IFC images of cells immediately after labeling with DiI (0h) and 22 h and 46 h post-labeling. DiI fluorescence is distributed throughout the cells, at the membrane and in intracellular compartments. EVs were collected in the conditioned medium between 22 and 46 h after DiI staining. **(B)** Representative IFC images of unlabelled (top) and DiI-labelled N2a cells (bottom). Images taken in the brightfield and DiI channels are shown. **(C)** Representative histogram of DiI cell intensity for unlabelled (left) and labelled cells (right). The bar graph on the right shows that the labeling procedure results in over 95% of cells that are DiI-positive up to 46 h following cell staining. **(D)** The median DiI intensity of the cells is not significantly affected after 22 h in culture but is decreased at 46 h after cell staining. Values are means +/- SD of 4 experiments analyzed by one-way ANOVA with Dunnet’s multiple comparisons test. \**p*< 0.05. **(E)** The median DiI fluorescence of EVs detected in the cleared conditioned medium does not significantly change at 24, 28 and 48h after cell staining (corresponding to 2, 4 and 24h collection). Values are means +/- SD of 4 experiments analyzed by one-way ANOVA with Dunnet’s multiple comparisons test. *p*>0.5.

The conditioned medium containing EVs secreted by DiI-stained cells was cleared from cell debris and apoptotic bodies by centrifugation at 2,000 x *g* for 10 min (cleared conditioned medium) and directly analyzed by IFC or further separated by ultracentrifugation or ultrafiltration-size exclusion chromatography (Fig. 1). IFC has recently emerged as a powerful approach for single particle analysis of fluorescently-labelled EVs (Gorgens et al., 2019; Lannigan and Erdbruegger, 2017). EVs secreted by DiI-stained cells were amenable to single particle analysis by IFC. The majority of the low scatter events detected in the cleared conditioned medium (82.7 ± 15.0 %) were DiI-positive (Fig 3A). Moreover, 98.8 ± 0.1 % of the DiI-positive events were detected as single objects by IFC. DiI-positive events were highly sensitive to lysis by the detergent NP-40, suggesting they were *bona fide* EV particles (Lannigan and Erdbruegger, 2017; Thery et al., 2018). On the contrary, DiI-negative events were not significantly affected by incubation with NP-40, suggesting they were not membrane-enclosed particles (Fig. 3B). The number of DiI-positive events detected in control samples such as buffer, unconditioned medium or medium conditioned by unstained cells was two orders of magnitude lower than the number of DiI-positive particles in the cleared conditioned medium from stained cells (Fig. 3C). Altogether, these data suggest that the conditioned medium of DiI-stained cells contains fluorescent objects that fulfill the definition of EVs and that can be efficiently detected by IFC. Importantly, the number of DiI positive particles detected by IFC in the cleared conditioned medium of stained cells correlated significantly well with the DiI fluorescence detected by spectrofluorometry (Fig. 3D), which could therefore be used as an alternative approach for the fast screening of changes in EV secretion by fluorescently labeled cells.

**Fig. 3 –.**
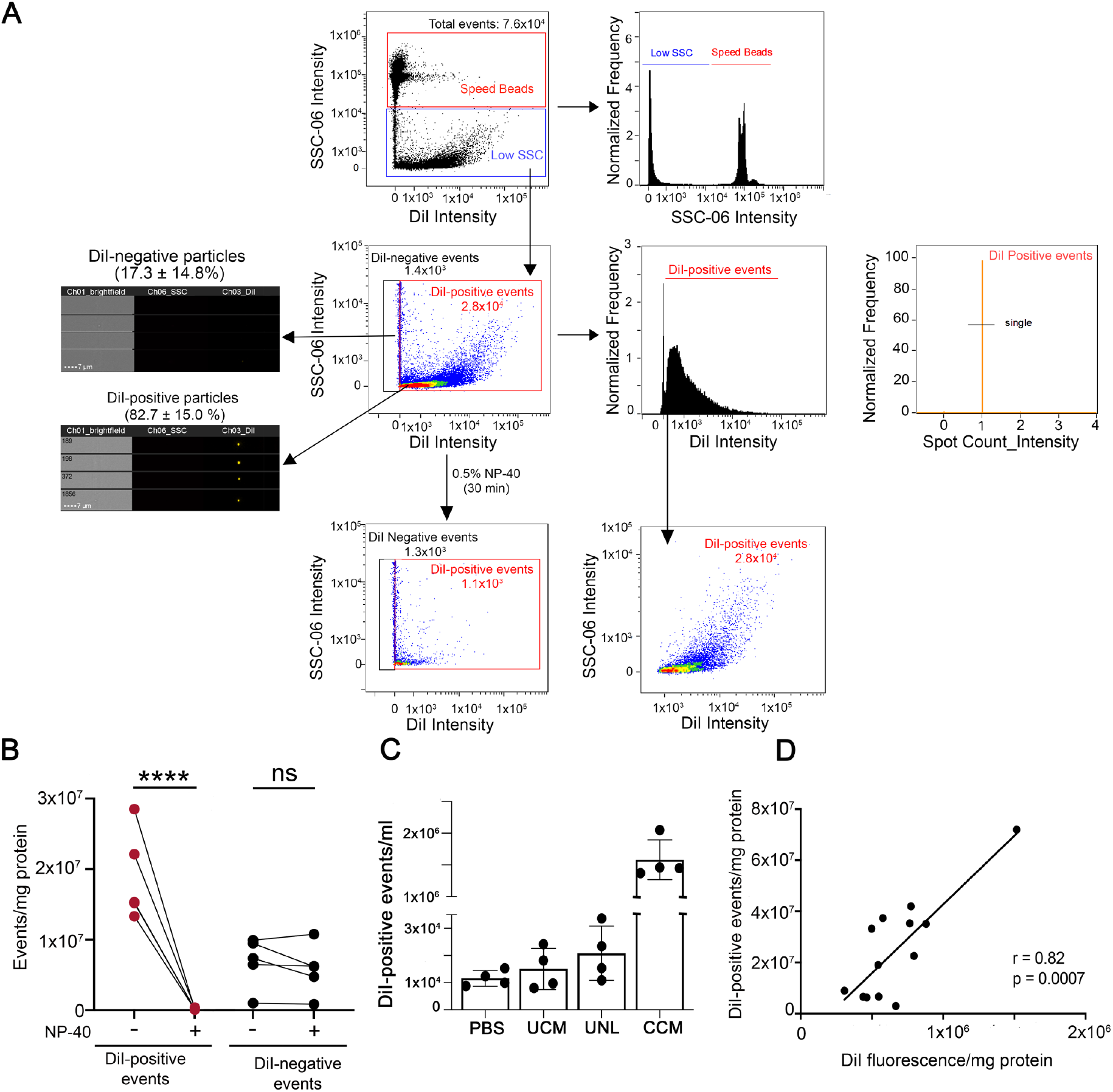
IFC allows the detection of EVs released in the cleared conditioned medium by DiI-labeled cells. **(A)** Gating strategy for EV analysis. First samples were gated on intensity in the scatter channel (Ch06 (SSC) to remove any remaining SpeedBeads from the analysis. A representative density dot plot of the low scatter population is included together with images taken in the brightfield, side scatter (SSC) and DiI channels of DiI-negative and DiI positive events. The majority of events are DiI-positive and disappear (are lysed) upon sample incubation with the detergent NP-40 (lower dot plot), demonstrating that they are membrane-enclosed particles. The histogram of DiI intensity on the right determines the population of DiI-positive events. To ensure the analysis of only 1 EV in each image, a mask was created to identify DiI intensity (Intensity (M03, Ch03_DiI, 50-4095)). Using this mask, we developed a spot count feature (Spot Count_Intensity (M03,Ch03_DiI, 50-4095)_4) to gate on images that had no more than 1 DiI spot (single). **(B)** Quantification of the effects of NP-40 on DiI-positive and DiI-negative particles in 4 independent experiments. While DiI-positive particles are highly sensitive to lysis by NP-40, DiI-negative particles are not, indicating they are not EVs. Data were analyzed by one-way ANOVA with Sidak’s multiple comparisons test. \*\*\*\**p*< 0.001, ns= not significant. **(C)** Quantification of DiI-positive events detected in buffer (PBS), unconditioned medium (UCM), medium conditioned by unlabeled cells (UNL) and cleared conditioned medium (CCM) from DiI-stained cells in 4 independent experiments. All samples were subjected to the same procedures, including the centrifugation at 2000 x *g* required to obtain the CCM and were measured for the same time. **(D)** Pearson’s correlation analysis between the number of DiI-positive events detected by IFC and DiI fluorescence measured by spectrofluorometry in the cleared conditioned medium.

In the experiments described above, we used a 488 nm laser to detect labelled EVs, and a 785 nm laser for side scatter (SSC) detection. Although it was proposed that the use of a 405 nm laser might have advantages over the 785nm laser (Lannigan and Erdbruegger, 2017), in our experiments we did not find that to be the case, as the 405 nm laser only increased the detection of background noise (Supplementary Fig. 3).

### Cell culture conditions

Careful consideration should be given to the medium in which the cells are incubated during EV collection to prevent potential confounding effects arising from the presence of serum-derived EVs and serum nanoparticle components (Coumans et al., 2017). As others before us (Lehrich et al., 2021), we found that even established protocols that involve ultracentrifugation at 100,000 x g for 16 h to deplete EVs from serum (EV-depleted serum) result in a significant amount of EVs still remaining in the serum (Supplementary Fig. 4). Therefore, unless otherwise specified, in our studies, EVs secreted by N2a cells were collected in medium that did not contain serum. To prevent possible effects of serum depletion on cells that could potentially affect EV secretion (Aswad et al., 2016; Eitan et al., 2015; Gardiner et al., 2016) the culture medium used for the collection period was carefully selected to allow optimal cell survival and to minimize the effects of the switch to nutrient-poor medium on EV secretion, including those caused by changes in autophagy. To this end, we compared cells grown in regular culture medium (DMEM:OptiMEM, 1:1 with sodium pyruvate and L-glutamine supplements, as indicated in Materials and Methods) containing 10% EV-depleted fetal bovine serum (EVd), with cells grown in DMEM:OptiMEM (1:1) with sodium pyruvate and L-glutamine supplements alone (Opti) or supplemented with N-2 or B27 supplements. N2a cell viability was not significantly affected by the lack of serum in the medium (Supplementary Figs. 5 A,B). However, only cells grown in DMEM:OptiMEM + N-2 supplement (N-2 medium) displayed similar metabolic activity (Supplementary Fig. 5B), autophagic activity (Supplementary Fig. 5 C), and secretion of EVs (Supplementary Fig. 6) compared to cells grown in serum-containing medium. Therefore, N-2 medium was used for EV collection in all experiments. All parameters described here, including the use of alternative culture media should be tested and validated for each different cell type.

### Comparative analysis of EVs in the cleared conditioned medium and upon separation by ultracentrifugation or ultrafiltration-size exclusion chromatography

Next, we sought to determine whether the fraction used for EV analysis and the method of EV separation affect EV recovery or skews the analysis towards specific subpopulations, based on size and EV markers. In particular, we compared data obtained from the analysis by IFC of EVs in the cleared conditioned medium, with data obtained after EV separation with established and widely used methods, such as ultracentrifugation at 100,000 x *g* (UC) and ultrafiltration-size exclusion chromatography (SEC), following the protocol illustrated in Fig. 1. Transmission electron microscopy of EVs separated by UC (Pellet_100K_) or SEC (SEC_peak_) demonstrated, as expected, the presence of characteristic cup-shaped EVs in both preparations (Fig. 4A). The Pellet_100K_ also contained several EV aggregates, a common technical artifact of UC procedures (Linares et al., 2015). The immunoblots in Fig. 4A show quality controls for the cleared conditioned medium, the Pellet_100K_ and the SEC_peak_, including the presence of the EV marker CD9 and the absence of calnexin, which indicates lack of contamination from apoptotic bodies and cell debris. The procedure of SEC successfully separated EVs from soluble proteins, and DiI fluorescence was exclusively present in the EVs fraction, as expected for EVs labeled incell (Fig. 4A). Both UC and SEC procedures for EV separation resulted in very low yields, as indicated by the significant decrease in the number of particles detected by IFC and nanoparticle tracking analysis (NTA) after UC and SEC, compared to the original unprocessed cleared conditioned medium (table in Fig. 4B). Compared to the latter, both methods indicate that less than 10% of EVs were recovered in the Pellet_100K_ after UC, mainly because a significant proportion of EVs remained in the supernatant. Similarly, the number of EVs recovered by SEC (SEC peak) corresponded to only 12.6 ± 0.99 % (by IFC) and 20.12 ± 6.23 % (by NTA) of the total EVs in the cleared conditioned medium. However, the EV particle size distribution as well as mode size assessed by NTA were not significantly affected by the sample or isolation technique used (Fig. 4C).

**Fig. 4.**
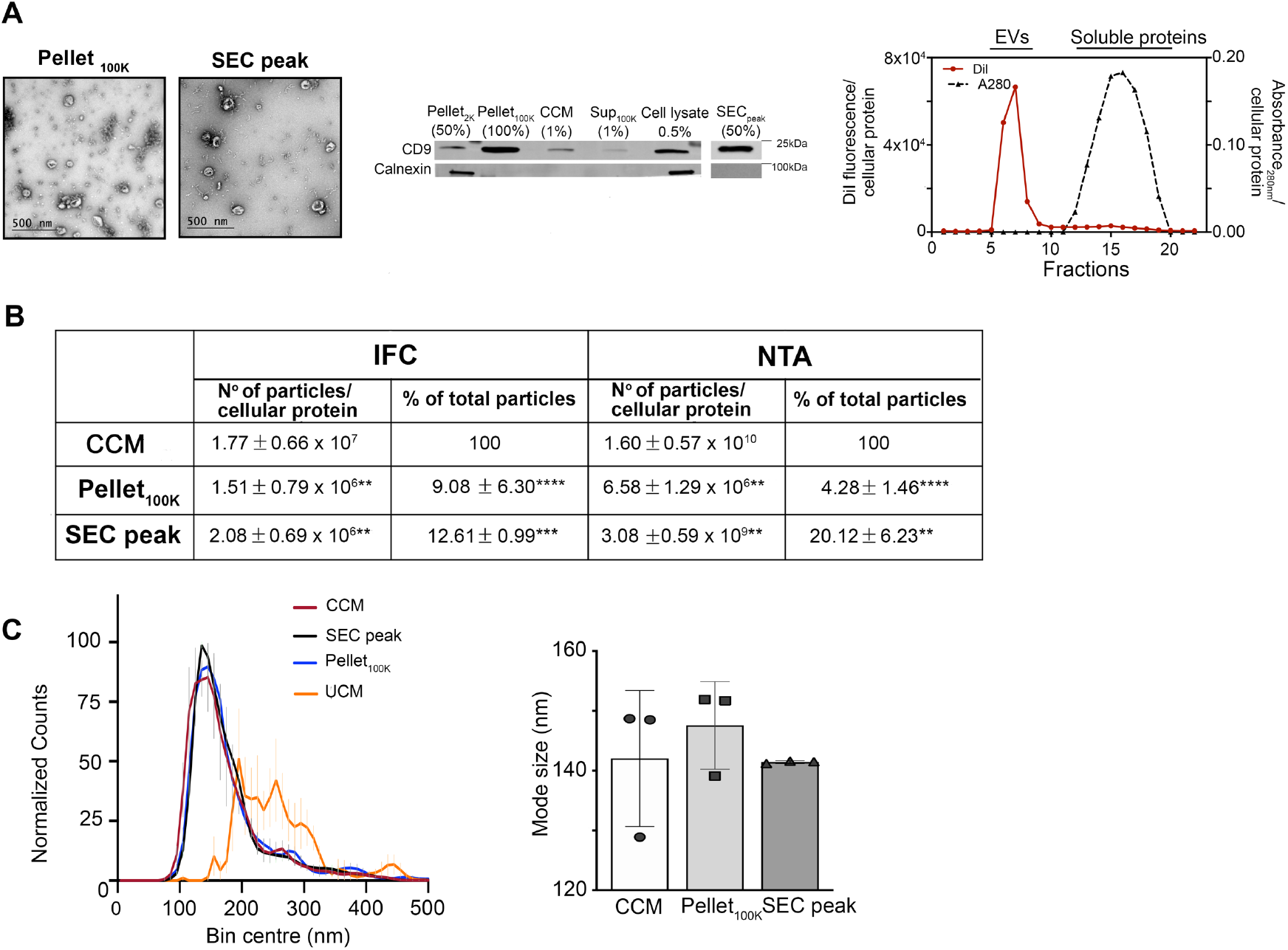
Characterization of EVs separated by sequential ultracentrifugation or ultrafiltration-size exclusion chromatography. EVs were separated by UC and SEC as indicated in Fig. 1. **(A)** Electron micrographs of Pellet_100K_ and SEC_peak_ fractions show characteristic cup-shaped EV particles. The immunoblot shows the presence of the EV marker CD9 and the absence of calnexin in all fractions obtained from the conditioned medium after the initial centrifugation at 2,000 x *g*, indicating the fractions are not contaminated by apoptotic bodies or cell debris. Numbers in parenthesis indicate the fraction for each sample that was loaded in the gel. A representative chromatogram of EV separation by SEC is shown on the right. **(B)** Summary of the average number of particles present in each fraction (normalized for cellular protein content) and percentage relative to the total number of particles detected in the cleared conditioned medium (CCM). Particle numbers were determined by imaging flow cytometry (IFC) and nanoparticle tracking analysis (NTA). Values are means ± SD of 6 (IFC) or 3 (NTA) independent experiments. Paired t-test was applied to compare the Pellet_100K_ or the SEC peak to the cleared conditioned medium. \*\*\*\**p*<0.0001, \*\*\**p*<0.001, \*\**p*<0.01 **(C)** Particle size distribution profiles of EVs in the cleared conditioned medium and after isolation by UC or SEC, as detected by NTA. Each profile was obtained using the mean values from 3 experiments. Vertical lines are SD for each size bin. The graph on the right shows the mode size for each indicated fraction. Particle size is not significantly different among fractions. Data are mean values ± SD of 3 independent experiments. \*\*\**p*<0.001. CCM = cleared conditioned medium, UCM = Unconditioned medium.

Given the remarkable loss of EVs resulting from UC and SEC separation procedures, we investigated whether the EVs recovered by UC are representative of the total EV population present in the unprocessed cleared conditioned medium, or of distinct EV subpopulations that might preferentially pellet at 100,000 x *g*. IFC facilitates the detection of EV subpopulations in a heterogeneous sample (Gorgens et al., 2019). To label distinct EV subpopulations, we incubated the cleared conditioned medium with anti-CD9 antibodies and Annexin V (which binds to phosphatidylserine, PS) prior to EV separation by UC. Based on these markers, IFC analysis of DiI-labeled particles showed the presence of four different EV populations in the cleared conditioned medium and in the Pellet_100K_ (Fig. 5A): DiI^+^/CD9^-^/Annexin^-^-EVs represented the larger population (~60% of total), followed by DiI^+^/CD9^+^/Annexin^-^-EVs (~20%), with the remaining ~20% divided between DiI^+^/CD9^-^/Annexin^+^-EVs and DiI^+^/CD9^+^/Annexin^+^-EVs. These four populations were equally represented in the cleared conditioned medium and in the Pellet_100K_ (Fig. 5C and Supplementary Fig. 7).

**Fig. 5.**
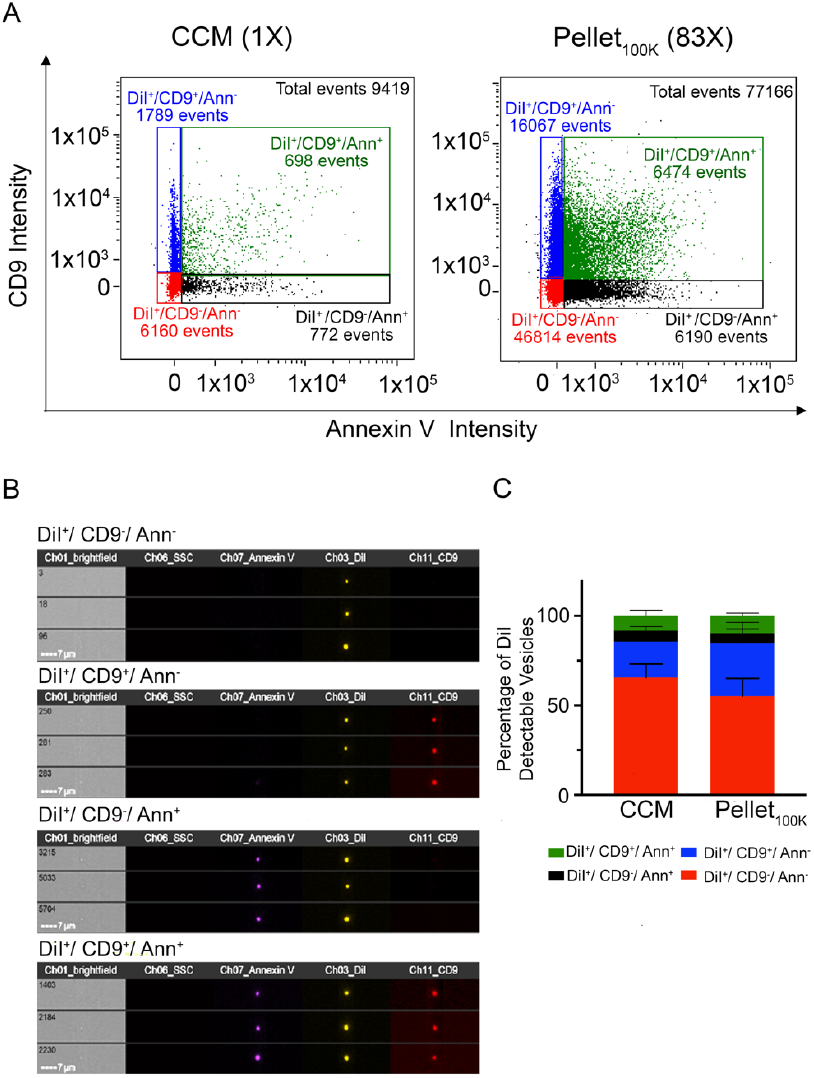
Comparison of markers distribution between the cleared conditioned medium and the Pellet_100K_. **(A)** Representative IFC dot plots for EVs labeled *in vivo* with DiI and *in vitro* with anti-CD9 antibodies and/or Annexin V. Analysis were performed in cultured conditioned medium (CCM) and the pellets that resulted from UC separation (Pellet_100K_). Numbers in parenthesis indicate the concentration factor in each sample. All samples were analyzed for the same time (10 min) in the same conditions. **(B)** Representative IFC images of particles in each subpopulation. **(C)** Quantification of the relative abundance of differentially labelled EV subpopulations in the cleared conditioned medium and in the Pellet_100K_. Data are mean percent values of detectable vesicles ± SD of 4 independent experiments. No significant differences between groups were found.

### Direct analysis of EVs in the cleared conditioned medium after cell treatment with bafilomycin

To assess the suitability of the direct analysis of the cleared conditioned medium by IFC to study the regulation of EV secretion, we used bafilomycin (Baf), a compound that blocks autophagic flux and consequently increases EV secretion (Baixauli et al., 2014; Cashikar and Hanson, 2019; Savina et al., 2003). We selected the minimum concentration of Baf required to achieve maximum autophagic blockade in N2a cells (Supplementary Fig. 8). EVs were collected from DiI-labeled N2a cells treated with Baf (150 nM) or vehicle for 4 h and analyzed by IFC directly in the cleared conditioned medium, or after separation by UC, in the Pellets_100K_. Representative dot plots are shown in Supplementary Fig 9. Analysis of the cleared conditioned medium, but not the Pellet_100K_, revealed the expected increase in the number of EVs secreted upon cell treatment with Baf (Fig. 6A). Moreover, only the analysis performed in the cleared conditioned medium allowed to detect small but significant changes in the relative abundance of different EV subpopulations, with an increase in DiI^+^/CD9^-^/Ann^-^ EVs (Fig. 6B and Supplementary Fig. 10).

**Fig. 6.**
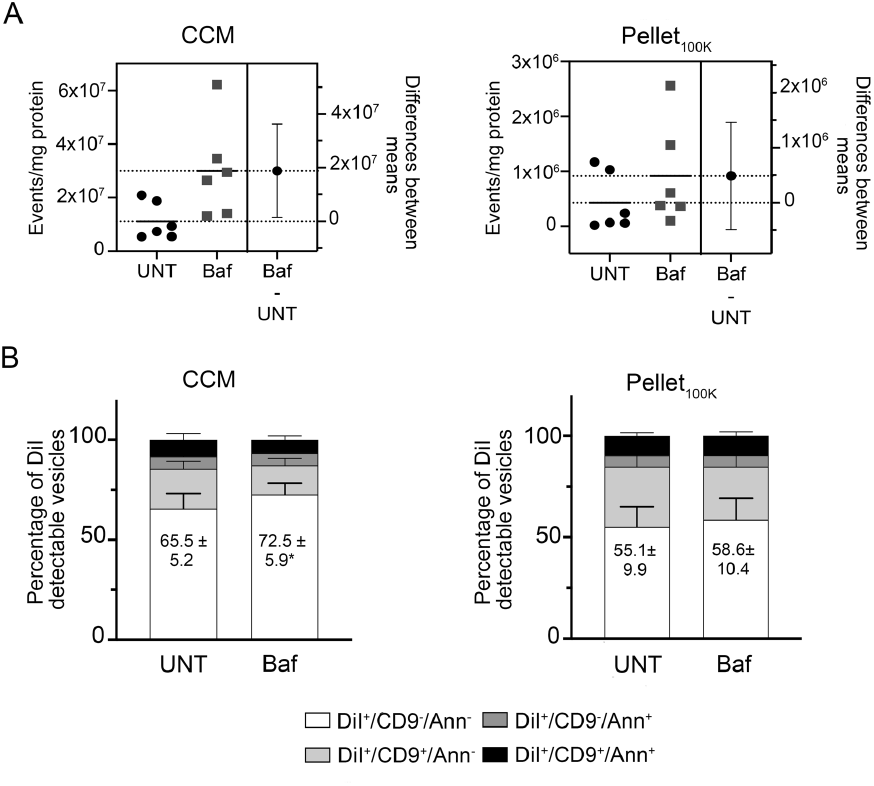
Effect of bafilomycin on EV release measured in the cleared conditioned medium and in the Pellet_100K_. EVs were collected from DiI-labeled N2a cells treated with Baf (150 nM) or vehicle (UNT) for 4 h and analyzed in the cleared conditioned medium (CCM) or after isolation by UC. The cleared conditioned media were adjusted based on protein content of donor cells before isolation by UC and analysis by IFC. All samples were counted for the same time (10 min). **(A)** Estimation plots for the total number of DiI-positive EVs in the cleared conditioned medium and in the Pellet_100K_, respectively. The left panel in each plot shows data for individual experiments and their mean. The right panel shows the effect size (difference between means) and its 95% confidence interval. **(B)** Cleared conditioned medium and Pellet_100K_ preparations were labeled *in vitro* using anti-CD9 antibodies and Annexin V. Graphs show the relative abundance of differentially labelled EV subpopulations in each fraction. Data are mean percent values ± SD for three independent experiments. Significance determined by unpaired *t*-test; \**p*<0.05.

## Discussion

### EVs labeling

In order to use IFC, uniform and bright fluorescent labeling of EVs is required. Our simple and versatile protocol of *in cell* labeling with the lipophilic dye DiI labeled both plasma and intracellular membranes of N2a cells. Although a decrease in cell labeling was observed at the end of the experiment, it was not reflected by a detectable decrease of EV labeling, indicating that pre-labeling of cells with DiI results in the stable incorporation of the dye into EVs, and allows for reliable EV detection by IFC in a timeframe that is suitable for most cell-based studies. However, we cannot exclude the possibility that vesicles harboring low levels of DiI would be missed using this protocol for EV labeling and detection.

Using NTA we identified EVs of varying size, suggesting that DiI labels EV of different cellular origin. We found a strong positive correlation between the number of DiI-labeled objects detected by IFC in the cleared conditioned medium and the DiI fluorescence assessed by spectrofluorometry, suggesting that measurements of fluorescence in the cleared conditioned medium from DiI-stained cells would provide a rapid estimation of EV secretion without the need of further processing. We obtained comparable results labeling the cells with DiD (not shown).

Many current strategies to label EVs released by cultured cells are based on labeling EVs *in vitro* after separation, or labeling parental cells, which results in the release of fluorescent EVs. Methods to label EVs *in vitro* have used many compounds including amine reactive dyes (carboxyfluorescein succinimidyl ester CFSE) (Morales-Kastresana et al., 2017; Pospichalova et al., 2015), lipophilic dyes (PKH dyes, DiI, DiD, DiR, etc.) (Grange et al., 2014; Morales-Kastresana et al., 2017; Nicola et al., 2009; Pospichalova et al., 2015; Puzar Dominkus et al., 2018; Tamura et al., 2016; van der Vlist et al., 2012; Wiklander et al., 2015) and Mem dyes (Shimomura et al., 2021) among others (Panagopoulou et al., 2020). Labeling of EVs *in vitro* suffers from the limitations inherent to the method employed for EV separation prior to labeling (discussed below) in addition to specific complications that arise during the labeling procedure, depending on the fluorophore employed. Dye aggregation has been reported during the labeling process using PKH dyes and DiI, which makes EV labeling *in vitro* with those lipid dyes unreliable, unless rigorous controls are used to discriminate between labelled EVs and non-EVfluorescent particles (micelles, nanoparticles or aggregated dye) (Lai et al., 2015; Morales-Kastresana et al., 2017; Puzar Dominkus et al., 2018; Russell et al., 2019). In addition, labeling EVs with PKH may cause a size shift towards larger size vesicles potentially through PKH nanoparticles fusion/aggregation (Dehghani et al., 2020). Moreover, the procedure requires dye elimination after labeling and the methodology used for that purpose also play a significant role in the final product (Puzar Dominkus et al., 2018). Labeling *in vitro* with CFSE prevents artifacts due to dye aggregation and does not affect the size of EVs (Dehghani et al., 2020), but although some studies suggested that removal of the unbound CFSE is not required (Pospichalova et al., 2015), others detected increased fluorescence in the background noise, possibly as a result of spontaneous hydrolysis of free dye (Morales-Kastresana et al., 2017). Similarly, no large aggregation and no significant change of the apparent size of EVs were observed with the Mem dyes, although excess dye needs to be eliminated after labeling (Shimomura et al., 2021).

Labeling of parental cells with different compounds such as CFSE (Ender et al., 2019), PKH dyes (Dabrowska et al., 2018; van der Vlist et al., 2012), DiI and DiD (Grange et al., 2014) have produced mixed results. CFSE may be toxic to the cells at the concentration required to label EVs; vesicles may not harbor enough CFSE fluorescence to be detected by flow cytometry (van der Vlist et al., 2012) and only a small sub-fraction of the total EV population secreted, corresponding to the pellet after 10,000x *g* centrifugation, is labeled (Ender et al., 2019).

In our method, even though labeling was performed at presumably saturating concentrations of DiI (based on manufacturer’s instruction) we did not observe cytotoxicity due to cell over-labeling as reported for other chromophores (Samlowski et al., 1991). Because the cells are labeled in bulk, before seeding them in culture, all experimental groups are labelled equally. Moreover, our method offers a simple solution to determine if different cell types or different variants (e,g. wild type and mutant cells) are labeled differently under similar conditions.

Another strategy to label EVs *in vivo* involves expression of fluorescent reporters fused to EV-specific proteins markers (Mittelbrunn et al., 2011; Piao et al., 2018; Verweij et al., 2018) or carrying farnesylation or palmitoylation consensus sequences (Lai et al., 2015). In the first scenario, there have been reports on the low proportion of EVs labeled (Wiklander et al., 2015) as well as the fact that labeling may be restricted to selective subpopulations of EVs, limiting the observations to only a few subtypes of EVs. The expression of membrane-bound fluorescent proteins overcomes the latter limitation, however the fluorescence intensity of EVs depends on protein expression level on the EV membrane, and it has been suggested that the expression of fluorescent proteins may affect EV properties and cargos. (Chuo et al., 2018). In summary, most current approaches for EVs labeling present limitations and there is a recognized need for better pan-EV dyes (Russell et al., 2019). Recently, metabolic labeling of EVs with azido sugars- or phospholipid-based biorthogonal conjugation have been reported, mostly to study *in vivo* biodistribution of EVs (Lee et al., 2018; Zhang et al., 2018). Although these strategies would label all EV populations independently of their cellular origin and would not influence EV proteins nor affect the structural integrity of EVs, the labeling protocols still require the separation of the excess fluorescent dye following the *in vitro* click chemistry reaction.

While optimization of techniques for labeling EVs *in vitro* is important for the analysis of EVs in biological samples and/or to accurately follow EV biodistribution *in vivo*, we report here a simple method to label EVs released by cultured cells that can be reliably used for the study of EV secretion regulation.

### EVs separation and analysis

EVs separated by UC and SEC were analyzed in parallel with EVs present in the cleared conditioned medium before separation, using IFC and NTA in a complementary approach. Both NTA and IFC provide the concentration of particles in a sample. NTA works by relating Brownian motion to particle size to determine number and size (Gardiner et al., 2013), while IFC works by detecting fluorescent energy from dyes or probes bound to EVs. IFC has addressed many of the limitations of traditional flow cytometry for measuring EVs with the added advantage of imagery confirmation (Erdbrugger and Lannigan, 2016; Lannigan and Erdbruegger, 2017; Mastoridis et al., 2018), but does have its own limitations as well (necessity of fluorescent signal for small particle detection). Similar data would be expected from any high-resolution flow cytometry platform that can resolve extracellular vesicles.

Our analysis confirmed several issues associated with the separation of EVs by UC or SEC. In agreement with previous reports (Mastoridis et al., 2018), NTA detected significantly higher numbers of particles than IFC in the same samples, because it also detect non-EV particles such as large protein aggregates and dus (Filipe et al., 2010), as well as EVs with low DiI fluorescence that might escape detection by IFC. Despite these differences, both IFC and NTA indicated very poor EV yield using UC and SEC as shown before (Brennan et al., 2020; Livshits et al., 2015). Although NTA analysis agreed with earlier indications that UC causes more significant reduction of particle yield than SEC (Lobb et al., 2015), analysis by IFC showed no significant differences in total number of EVs isolated by UC and SEC and there was no difference in the levels of CD9 detected in EVs separated by either method. An additional disadvantage of EV separation by UC is the artificial aggregation and/or fragmentation of EVs, which might lead to artifacts during single particle analysis and flow cytometry analysis (Erdbrugger and Lannigan, 2016) and may mask antigens on the EV surface, thus complicating phenotypic analysis of EVs based on markers (Linares et al., 2015). Furthermore, aggregation and co-sedimentation of free proteins with the EV pellet may cause EV contamination (Lamparski et al., 2002) that may remain undetected depending of the method employed for EVs analysis. Despite the significant loss of EVs upon isolation by UC, the proportion of DiI-detectable vesicles in each of the four EV populations analyzed in our studies was unchanged, although we cannot exclude that other EV subpopulations might be affected by UC.

Our studies demonstrated for the first time that direct analysis of the cleared conditioned medium may allow to detect changes in EV secretion that might be otherwise missed if EVs are isolated by UC, at least when the changes are of a relatively small magnitude. In our experiments with bafilomycin, the expected increase of EV release was detected by IFC analysis of the cleared conditioned medium, while analysis of the pellet after UC provided non-significant differences in EV secretion. Moreover, the relative abundance of EV subpopulations in the cleared conditioned medium revealed subtle drug-induced changes that could not be detected in the Pellet_100K_. Therefore, all considered, we propose that for the study of EV secretion, direct analysis of EVs in the cleared conditioned medium is preferable to the analysis in the pellet 100K, unless, of course, maximum EV concentration or separation from soluble components in the medium (proteins and other factors) are required for downstream applications. Moreover, analysis of the cleared conditioned media should be an integrated component of any EV study so as to reduce the chance of losing valuable data due to technical artifacts.

In conclusion:

- We presented a protocol that makes use of a unique combination of in cell-labeling and EV analysis by IFC directly in the cleared conditioned medium.
- This protocol offers higher EV yields and prevents selective loss of EV subpopulations that can occur with SEC- and UC-based protocols, which makes it more suitable than the latter for the characterization of EV subpopulation secreted by cells in the conditioned medium.
- Altogether, the data presented here indicate that cell pre-labeling with DiI, coupled with IFC analysis of fluorescent EV particles directly in the cleared conditioned medium represents a powerful and convenient method to study EV secretion *in vitro*, with minimal sample handling and loss.

## Abbreviations

Baf: Bafilomycin A1;
BCA: Bicinchoninic acid;
EVs: Extracellular vesicles;
CCM: cleared conditioned medium;
FBS: Fetal bovine serum;
IFC: Image Flow Cytometry;
LC3: Microtubule-associated protein light chain 3;
NTA: Nanoparticle tracking analysis;
PBS: Phosphate buffered saline;
PS: Phosphatidylserine;
SEC: Size exclusion chromatography;
TEM: Transmission electron microscopy;
UC: Differential ultracentrifugation;
UCM: Unconditioned media.

## ACKNOWLEDGMENTS

This work was supported by grants from the Alberta Innovates/Alberta Prion Research Institute (201800001/RES0039430) and Synergies in Alzheimer Disease Research (SynAD) to S. Sipione and E. Posse de Chaves, Brain Canada (RES0030547), GlycoNet (RES0050801) and NSERC (RES0029350) to S. Sipione, and from the Alzheimer Society of Alberta and Northwestern Territories/ Alberta Innovates/Alberta Prion Research Institute (201600003/RES0029396) to E. Posse de Chaves. A.M. Rieger is an ISAC SRL Emerging Leader (2017-2021). Imaging Flow Cytometry experiments were performed at the University of Alberta Faculty of Medicine & Dentistry Flow Cytometry Facility, RRID:SCR_019195, which receives financial support from the Faculty of Medicine & Dentistry and Canada Foundation for Innovation (CFI) awards to contributing investigators. We thank Ms. S. Samuelson for technical support and Mr. Rohan Aananth and Dr. Qian Wang for performing preliminary experiments on cell labeling and antibodies titration.

## References

Andjus, P., M. Kosanovic, K. Milicevic, M. Gautam, S.J. Vainio, D. Jagecic, E.N. Kozlova, A. Pivoriunas, J.C. Chachques, M. Sakaj, G. Brunello, D. Mitrecic, and B. Zavan. 2020. Extracellular Vesicles as Innovative Tool for Diagnosis, Regeneration and Protection against Neurological Damage. Int J Mol Sci. 21.

Aswad, H., A. Jalabert, and S. Rome. 2016. Depleting extracellular vesicles from fetal bovine serum alters proliferation and differentiation of skeletal muscle cells in vitro. BMC Biotechnol. 16:32.

Baixauli, F., C. Lopez-Otin, and M. Mittelbrunn. 2014. Exosomes and autophagy: coordinated mechanisms for the maintenance of cellular fitness. Front Immunol. 5:403.

Born, L.J., J.W. Harmon, and S.M. Jay. 2020. Therapeutic potential of extracellular vesicle-associated long noncoding RNA. Bioeng Transl Med. 5:e10172.

Brennan, K., K. Martin, S.P. FitzGerald, J. O’Sullivan, Y. Wu, A. Blanco, C. Richardson, and M.M. Mc Gee. 2020. A comparison of methods for the isolation and separation of extracellular vesicles from protein and lipid particles in human serum. Sci Rep. 10:1039.

Cashikar, A.G., and P.I. Hanson. 2019. A cell-based assay for CD63-containing extracellular vesicles. PLOS ONE. 14:e0220007.

Chuo, S.T., J.C. Chien, and C.P. Lai. 2018. Imaging extracellular vesicles: current and emerging methods. J Biomed Sci. 25:91.

Coumans, F.A.W., A.R. Brisson, E.I. Buzas, F. Dignat-George, E.E.E. Drees, S. El-Andaloussi, C. Emanueli, A. Gasecka, A. Hendrix, A.F. Hill, R. Lacroix, Y. Lee, T.G. van Leeuwen, N. Mackman, I. Mager, J.P. Nolan, E. van der Pol, D.M. Pegtel, S. Sahoo, P.R.M. Siljander, G. Sturk, O. de Wever, and R. Nieuwland. 2017. Methodological Guidelines to Study Extracellular Vesicles. Circ Res. 120:1632–1648.

Dabrowska, S., A. Del Fattore, E. Karnas, M. Frontczak-Baniewicz, H. Kozlowska, M. Muraca, M. Janowski, and B. Lukomska. 2018. Imaging of extracellular vesicles derived from human bone marrow mesenchymal stem cells using fluorescent and magnetic labels. International Journal of Nanomedicine. Volume 13:1653–1664.

Dehghani, M., S.M. Gulvin, J. Flax, and T.R. Gaborski. 2020. Systematic Evaluation of PKH Labelling on Extracellular Vesicle Size by Nanoparticle Tracking Analysis. Scientific Reports. 10.

Eitan, E., S. Zhang, K.W. Witwer, and M.P. Mattson. 2015. Extracellular vesicle-depleted fetal bovine and human sera have reduced capacity to support cell growth. J Extracell Vesicles. 4:26373.

Ender, F., P. Zamzow, N. Von Bubnoff, and F. Gieseler. 2019. Detection and Quantification of Extracellular Vesicles via FACS: Membrane Labeling Matters! International Journal of Molecular Sciences. 21:291.

Erdbrugger, U., and J. Lannigan. 2016. Analytical challenges of extracellular vesicle detection: A comparison of different techniques. Cytometry A. 89:123–134.

Erdbrugger, U., C.K. Rudy, M.E. Etter, K.A. Dryden, M. Yeager, A.L. Klibanov, and J. Lannigan. 2014. Imaging flow cytometry elucidates limitations of microparticle analysis by conventional flow cytometry. Cytometry A. 85:756–770.

Gardiner, C., Y.J. Ferreira, R.A. Dragovic, C.W. Redman, and I.L. Sargent. 2013. Extracellular vesicle sizing and enumeration by nanoparticle tracking analysis. J Extracell Vesicles. 2.

Gardiner, C., D.D. Vizio, S. Sahoo, C. Théry, K.W. Witwer, M. Wauben, and A.F. Hill. 2016. Techniques used for the isolation and characterization of extracellular vesicles: results of a worldwide survey. Journal of Extracellular Vesicles. 5:32945.

Gorgens, A., M. Bremer, R. Ferrer-Tur, F. Murke, T. Tertel, P.A. Horn, S. Thalman, J.A. Welsh, C. Probst, C. Guerin, C.M. Boulanger, J.C. Jones, H. Hanenber, U. Erdbrugger, J. Lannigan, F.L. Ricklefs, S. El-Andaloussi, and B. Giebel. 2019. Optimisation of imaging flow cytometry for the analysis of single extracellular vesicles by using fluorescence-tagged vesicles as biological reference material. Journal of Extracellular Vesicles. 8.

Grange, C., M. Tapparo, S. Bruno, D. Chatterjee, P.J. Quesenberry, C. Tetta, and G. Camussi. 2014. Biodistribution of mesenchymal stem cell-derived extracellular vesicles in a model of acute kidney injury monitored by optical imaging. Int J Mol Med. 33:1055–1063.

Hu, T., J. Wolfram, and S. Srivastava. 2020. Extracellular Vesicles in Cancer Detection: Hopes and Hypes. Trends Cancer.

Inglis, H.C., A. Danesh, A. Shah, J. Lacroix, P.C. Spinella, and P.J. Norris. 2015. Techniques to improve detection and analysis of extracellular vesicles using flow cytometry. Cytom Part A. 87a:1052–1063.

Kim, I.A., J.Y. Hur, H.J. Kim, S.E. Lee, W.S. Kim, and K.Y. Lee. 2020. Liquid biopsy using extracellular vesicle-derived DNA in lung adenocarcinoma. J Pathol Transl Med.

Kostyushev, D., A. Kostyusheva, S. Brezgin, V. Smirnov, E. Volchkova, A. Lukashev, and V. Chulanov. 2020. Gene Editing by Extracellular Vesicles. Int J Mol Sci. 21.

Lai, C.P., E.Y. Kim, C.E. Badr, R. Weissleder, T.R. Mempel, B.A. Tannous, and X.O. Breakefield. 2015. Visualization and tracking of tumour extracellular vesicle delivery and RNA translation using multiplexed reporters. Nat Commun. 6:7029.

Lamparski, H.G., A. Metha-Damani, J.Y. Yao, S. Patel, D.H. Hsu, C. Ruegg, and J.B. Le Pecq. 2002. Production and characterization of clinical grade exosomes derived from dendritic cells. J Immunol Methods. 270:211–226.

Lannigan, J., and U. Erdbruegger. 2017. Imaging flow cytometry for the characterization of extracellular vesicles. Methods. 112:55–67.

Lee, T.S., Y. Kim, W. Zhang, I.H. Song, and C.-H. Tung. 2018. Facile metabolic glycan labeling strategy for exosome tracking. Biochimica et Biophysica Acta (BBA) - General Subjects. 1862:1091–1100.

Lehrich, B.M., Y. Liang, and M.S. Fiandaca. 2021. Foetal bovine serum influence on in vitro extracellular vesicle analyses. Journal of Extracellular Vesicles. 10.

Linares, R., S. Tan, C. Gounou, N. Arraud, and A.R. Brisson. 2015. High-speed centrifugation induces aggregation of extracellular vesicles. J Extracell Vesicles. 4:29509.

Livshits, M.A., E. Khomyakova, E.G. Evtushenko, V.N. Lazarev, N.A. Kulemin, S.E. Semina, E.V. Generozov, and V.M. Govorun. 2015. Isolation of exosomes by differential centrifugation: Theoretical analysis of a commonly used protocol. Sci Rep. 5:17319.

Lobb, R.J., M. Becker, S.W. Wen, C.S. Wong, A.P. Wiegmans, A. Leimgruber, and A. Moller. 2015. Optimized exosome isolation protocol for cell culture supernatant and human plasma. J Extracell Vesicles. 4:27031.

Margolis, L., and Y. Sadovsky. 2019. The biology of extracellular vesicles: The known unknowns. PLoS Biol. 17:e3000363.

Mastoridis, S., G.M. Bertolino, G. Whitehouse, F. Dazzi, A. Sanchez-Fueyo, and M. Martinez-Llordella. 2018. Multiparametric Analysis of Circulating Exosomes and Other Small Extracellular Vesicles by Advanced Imaging Flow Cytometry. Front Immunol. 9:1583.

Mathews, P.M., and E. Levy. 2019. Exosome Production Is Key to Neuronal Endosomal Pathway Integrity in Neurodegenerative Diseases. Front Neurosci. 13:1347.

Mehryab, F., S. Rabbani, S. Shahhosseini, F. Shekari, Y. Fatahi, H. Baharvand, and A. Haeri. 2020. Exosomes as a nextgeneration drug delivery system: An update on drug loading approaches, characterization, and clinical application challenges. Acta Biomater. 113:42–62.

Mittelbrunn, M., C. Gutierrez-Vazquez, C. Villarroya-Beltri, S. Gonzalez, F. Sanchez-Cabo, M.A. Gonzalez, A. Bernad, and F. Sanchez-Madrid. 2011. Unidirectional transfer of microRNA-loaded exosomes from T cells to antigen-presenting cells. Nat Commun. 2:282.

Morales-Kastresana, A., B. Telford, T.A. Musich, K. McKinnon, C. Clayborne, Z. Braig, A. Rosner, T. Demberg, D.C. Watson, T.S. Karpova, G.J. Freeman, R.H. DeKruyff, G.N. Pavlakis, M. Terabe, M. Robert-Guroff, J.A. Berzofsky, and J.C. Jones. 2017. Labeling Extracellular Vesicles for Nanoscale Flow Cytometry. Sci Rep. 7:1878.

Murao, A., M. Brenner, M. Aziz, and P. Wang. 2020. Exosomes in Sepsis. Front Immunol. 11:2140.

Nam, G.H., Y. Choi, G.B. Kim, S. Kim, S.A. Kim, and I.S. Kim. 2020. Emerging Prospects of Exosomes for Cancer Treatment: From Conventional Therapy to Immunotherapy. Adv Mater:e2002440.

Nicola, A.M., S. Frases, and A. Casadevall. 2009. Lipophilic Dye Staining of Cryptococcus neoformans Extracellular Vesicles and Capsule. Eukaryotic Cell. 8:1373–1380.

Panagopoulou, M.S., A.W. Wark, D.J.S. Birch, and C.D. Gregory. 2020. Phenotypic analysis of extracellular vesicles: a review on the applications of fluorescence. J Extracell Vesicles. 9:1710020.

Piao, Y.J., H.S. Kim, E.H. Hwang, J. Woo, M. Zhang, and W.K. Moon. 2018. Breast cancer cell-derived exosomes and macrophage polarization are associated with lymph node metastasis. Oncotarget. 9:7398–7410.

Pospichalova, V., J. Svoboda, Z. Dave, A. Kotrbova, K. Kaiser, D. Klemova, L. Ilkovics, A. Hampl, I. Crha, E. Jandakova, L. Minar, V. Weinberger, and V. Bryja. 2015. Simplified protocol for flow cytometry analysis of fluorescently labeled exosomes and microvesicles using dedicated flow cytometer. J Extracell Vesicles. 4:25530.

Puzar Dominkus, P., M. Stenovec, S. Sitar, E. Lasic, R. Zorec, A. Plemenitas, E. Zagar, M. Kreft, and M. Lenassi. 2018. PKH26 labeling of extracellular vesicles: Characterization and cellular internalization of contaminating PKH26 nanoparticles. Biochim Biophys Acta. 1860:1350–1361.

Raposo, G., and P.D. Stahl. 2019. Extracellular vesicles: a new communication paradigm? Nat Rev Mol Cell Biol. 20:509–510.

Royo, F., C. Théry, J.M. Falcón-Pérez, R. Nieuwland, and K.W. Witwer. 2020. Methods for Separation and Characterization of Extracellular Vesicles: Results of a Worldwide Survey Performed by the ISEV Rigor and Standardization Subcommittee. Cells. 9:1955.

Russell, A.E., A. Sneider, K.W. Witwer, P. Bergese, S.N. Bhattacharyya, A. Cocks, E. Cocucci, U. Erdbrugger, J.M. Falcon-Perez, D.W. Freeman, T.M. Gallagher, S. Hu, Y. Huang, S.M. Jay, S.I. Kano, G. Lavieu, A. Leszczynska, A.M. Llorente, Q. Lu, V. Mahairaki, D.C. Muth, N. Noren Hooten, M. Ostrowski, I. Prada, S. Sahoo, T.H. Schoyen, L. Sheng, D. Tesch, G. Van Niel, R.E. Vandenbroucke, F.J. Verweij, A.V. Villar, M. Wauben, A.M. Wehman, H. Yin, D.R.F. Carter, and P. Vader. 2019. Biological membranes in EV biogenesis, stability, uptake, and cargo transfer: an ISEV position paper arising from the ISEV membranes and EVs workshop. J Extracell Vesicles. 8:1684862.

Samlowski, W.E., B.A. Robertson, B.K. Draper, E. Prystas, and J.R. McGregor. 1991. Effects of supravital fluorochromes used to analyze the in vivo homing of murine lymphocytes on cellular function. J Immunol Methods. 144:101–115.

Sarko, D.K., and C.E. McKinney. 2017. Exosomes: Origins and Therapeutic Potential for Neurodegenerative Disease. Front Neurosci. 11:82.

Savina, A., M. Furlán, M. Vidal, and M.I. Colombo. 2003. Exosome Release Is Regulated by a Calcium-dependent Mechanism in K562 Cells. Journal of Biological Chemistry. 278:20083–20090.

Shimomura, T., R. Seino, K. Umezaki, A. Shimoda, T. Ezoe, M. Ishiyama, and K. Akiyoshi. 2021. New Lipophilic Fluorescent Dyes for Labeling Extracellular Vesicles: Characterization and Monitoring of Cellular Uptake. Bioconjugate Chemistry. 32:680–684.

Srivastava, A., S. Rathore, A. Munshi, and R. Ramesh. 2021. Extracellular Vesicles in Oncology: from Immune Suppression to Immunotherapy. The AAPS Journal. 23.

Stahl, P.D., and G. Raposo. 2019. Extracellular Vesicles: Exosomes and Microvesicles, Integrators of Homeostasis. Physiology (Bethesda). 34:169–177.

Tamura, R., S. Uemoto, and Y. Tabata. 2016. Immunosuppressive effect of mesenchymal stem cell-derived exosomes on a concanavalin A-induced liver injury model. Inflamm Regen. 36:26.

Thery, C., K.W. Witwer, E. Aikawa, M.J. Alcaraz, J.D. Anderson, R. Andriantsitohaina, A. Antoniou, T. Arab, F. Archer, G.K. Atkin-Smith, D.C. Ayre, J.M. Bach, D. Bachurski, H. Baharvand, L. Balaj, S. Baldacchino, N.N. Bauer, A.A. Baxter, M. Bebawy, C. Beckham, A. Bedina Zavec, A. Benmoussa, A.C. Berardi, P. Bergese, E. Bielska, C. Blenkiron, S. Bobis-Wozowicz, E. Boilard, W. Boireau, A. Bongiovanni, F.E. Borras, S. Bosch, C.M. Boulanger, X. Breakefield, A.M. Breglio, M.A. Brennan, D.R. Brigstock, A. Brisson, M.L. Broekman, J.F. Bromberg, P. Bryl-Gorecka, S. Buch, A.H. Buck, D. Burger, S. Busatto, D. Buschmann, B. Bussolati, E.I. Buzas, J.B. Byrd, G. Camussi, D.R. Carter, S. Caruso, L.W. Chamley, Y.T. Chang, C. Chen, S. Chen, L. Cheng, A.R. Chin, A. Clayton, S.P. Clerici, A. Cocks, E. Cocucci, R.J. Coffey, A. Cordeiro-da-Silva, Y. Couch, F.A. Coumans, B. Coyle, R. Crescitelli, M.F. Criado, C. D’Souza-Schorey, S. Das, A. Datta Chaudhuri, P. de Candia, E.F. De Santana, O. De Wever, H.A. Del Portillo, T. Demaret, S. Deville, A. Devitt, B. Dhondt, D. Di Vizio, L.C. Dieterich, V. Dolo, A.P. Dominguez Rubio, M. Dominici, M.R. Dourado, T.A. Driedonks, F.V. Duarte, H.M. Duncan, R.M. Eichenberger, K. Ekstrom, S. El Andaloussi, C. Elie-Caille, U. Erdbrugger, J.M. Falcon-Perez, F. Fatima, J.E. Fish, M. Flores-Bellver, A. Forsonits, A. Frelet-Barrand, et al. 2018. Minimal information for studies of extracellular vesicles 2018 (MISEV2018): a position statement of the International Society for Extracellular Vesicles and update of the MISEV2014 guidelines. J Extracell Vesicles. 7:1535750.

van der Vlist, E.J., E.N. Nolte-’t Hoen, W. Stoorvogel, G.J. Arkesteijn, and M.H. Wauben. 2012. Fluorescent labeling of nano-sized vesicles released by cells and subsequent quantitative and qualitative analysis by high-resolution flow cytometry. Nat Protoc. 7:1311–1326.

Verweij, F.J., M.P. Bebelman, C.R. Jimenez, J.J. Garcia-Vallejo, H. Janssen, J. Neefjes, J.C. Knol, R. De Goeij-De Haas, S.R. Piersma, S.R. Baglio, M. Verhage, J.M. Middeldorp, A. Zomer, J. Van Rheenen, M.G. Coppolino, I. Hurbain, G. Raposo, M.J. Smit, R.F.G. Toonen, G. Van Niel, and D.M. Pegtel. 2018. Quantifying exosome secretion from single cells reveals a modulatory role for GPCR signaling. Journal of Cell Biology. 217:1129–1142.

Wiklander, O.P.B., J.Z. Nordin, A. O’Loughlin, Y. Gustafsson, G. Corso, I. Mager, P. Vader, Y. Lee, H. Sork, Y. Seow, N. Heldring, L. Alvarez-Erviti, C.I.E. Smith, K. Le Blanc, P. Macchiarini, P. Jungebluth, M.J.A. Wood, and S. EL Andaloussi. 2015. Extracellular vesicle in vivo biodistribution is determined by cell source, route of administration and targeting. Journal of Extracellular Vesicles. 4.

Yan, H.C., T.T. Yu, J. Li, Y.Q. Qiao, L.C. Wang, T. Zhang, Q. Li, Y.H. Zhou, and D.W. Liu. 2020. The Delivery of Extracellular Vesicles Loaded in Biomaterial Scaffolds for Bone Regeneration. Front Bioeng Biotechnol. 8:1015.

Zhang, P., B. Dong, E. Zeng, F. Wang, Y. Jiang, D. Li, and D. Liu. 2018. In Vivo Tracking of Multiple Tumor Exosomes Labeled by Phospholipid-Based Bioorthogonal Conjugation. Analytical Chemistry. 90:11273–11279.

Zocchi, M.R., F. Tosetti, R. Benelli, and A. Poggi. 2020. Cancer Nanomedicine Special Issue Review Anticancer Drug Delivery with Nanoparticles: Extracellular Vesicles or Synthetic Nanobeads as Therapeutic Tools for Conventional Treatment or Immunotherapy. Cancers (Basel). 12.

