## Supplemental information for "A method to study extracellular vesicles secreted *in vitro* by cultured cells with minimum sample processing and extracellular vesicle loss"

### SUPPLEMENTARY MATERIALS AND METHODS

#### Preparation of EV collection media

Where indicated, EVs were collected in serum-free and phenol red-free medium supplemented with heat inactivated EV-depleted FBS (EVD complete medium). To prepare EV-depleted serum, heat inactivated FBS was transferred to 26.3mL thickwall polycarbonate bottles (Beckman Coulter, USA) and subjected to centrifugation at 100,000 x g for 16 h at 4°C using a Type 70 Ti Fixed-angle rotor (Beckman Coulter, USA) in an Optima L-70 ultracentrifuge (Beckman Coulter, USA) [427]. The resulting supernatant was filter-sterilized with a 0.22μm PVDF filter (Millipore, USA) and stored at -20°C until further use.

#### Evaluation of cell metabolic activity

Cell metabolic activity was evaluated using the MTT reduction assay (Promega). This assay is based on the conversion of MTT (3-(4,5-dimethylthiozolyl)-2,5-diphenyltetrazolium bromide) to a blue/purple formazan crystal by NADPH or NADH produced by dehydrogenase enzymes in metabolically active cells (Berridge and Tan, 1993). Cells were seeded in a 96-well plate at a density of $1.0x10^4$ cells/well and allowed to attach and grow for 48 h to a confluence of ~80% confluence, MTT assay was performed according to the manufacturer's instructions. Briefly, MTT labeling reagent was added (final concentration of 0.5mg/mL) to each well. The microplate was incubated at 37°C in 5% $CO_2$ for 4 h. Next, 100 μl of solubilization solution was added to each well. Following complete solubilization, the plate was read at 550nm using a SpectraMax® i3x multi-mode microplate reader (Molecular Devices, USA).

#### Iodixanol Flotation Gradient

Iodixanol (Optiprep™ Density Gradient Medium, Axis-Shield, Oslo, Norway) gradients were performed according to previously described protocols (Greening et al., 2015) with minor adjustments. Briefly, 5%, 10%, 20% and 40% iodixanol solutions were freshly prepared in serum-free medium (DMEM:OptiMEM 1:1) and rotated end-over-end at 4° C for 30 min. A discontinuous gradient was carefully layered by adding 3ml of 40%, 3ml of 20%, 3ml of 10%, and 2ml of 5%, solutions in a 16.8 ml open-top polyallomer tube (Beckman Coulter, Fullerton, California, USA). Gradient integrity was visually examined to ensure no mixing of layers occurred prior to sample addition. Cleared conditioned media were concentrated using Amicon Ultra-15 100K MWCO filters according to manufacturer's instructions. The resulting retentate (i.e., the volume retained on top of the filter) was brought up to 1ml with serum-free medium (DMEM:OptiMEM 1:1) and layered gently on the gradient at 4°C. Gradients were centrifuged using SW 41-Ti swinging bucket rotor (Beckman Coulter, Fullerton, California, USA) for 18 h at 100,000 × g. After centrifugation, 12 x 1ml fractions were collected from top to bottom. All steps were performed at 4°C. For immunoblotting, 10% of each fraction was precipitated with 4 volumes of ice-cold methanol overnight at −20°C. Following centrifugation at 16,000 × g for 15 min at 4°C, the protein pellet was washed once with methanol and centrifuged again. The final protein pellet was allowed to air dry for no longer than 5 min and was re-suspended in RIPA buffer with protease inhibitors. For dot-blot analysis, protein precipitation was not required and samples were loaded directly onto the membrane.

### Dot-blot analysis

Cells lysed in 1X NP-40 lysis buffer (20 mM Tris-HCl at pH 7.4, 1% NP-40 v/v, 1 mM EDTA, 1 mM EGTA, 50µM MG132, 1X protease/phosphatase inhibitor cocktail) were passed through a 27-G needle 10 times, incubated on ice for 30 min and then sonicated at intensity 2.0 for 10 s, prior to sample immobilization onto a 0.45µm pore size nitrocellulose membrane (Millipore, USA) using a 96-well Bio-Dot® apparatus (Bio-Rad, USA). EV samples were either sonicated at intensity 2.0 for 10 s or lysed with 0.01% NP-40. One microgram of proteins from each cell lysate was loaded into each well, in triplicates. Membranes were blocked for 1 h with Odyssey® blocking buffer (LiCor Biosciences, USA) or Intercept® blocking buffer (LiCor Biosciences, USA), and then incubated overnight at 4°C or for 2 h at r.t. with antibody anti-ALIX (AIP1) (1:250; BD Biosciences, 611621), diluted in Odyssey® or Intercept® blocking buffer. Membranes were then washed with TTBS (20mM Tris-HCl, 137mM NaCl, pH 7.4-7.6 containing 0.1% Tween-20) and incubated with fluorescent secondary antibody $\alpha$mouse IRDye®680 (1:5000). Finally, membranes were washed once with TBS-T and once more with TBS (20mM Tris-HCl, 137mM NaCl, pH 7.4-7.6) before scanning in an Odyssey® near-infrared scanner (LiCor Biosciences, USA). Signal was quantified using Odyssey® Application Software v3.0 (LiCor Biosciences, USA).

### SUPPLEMENTARY FIGURES

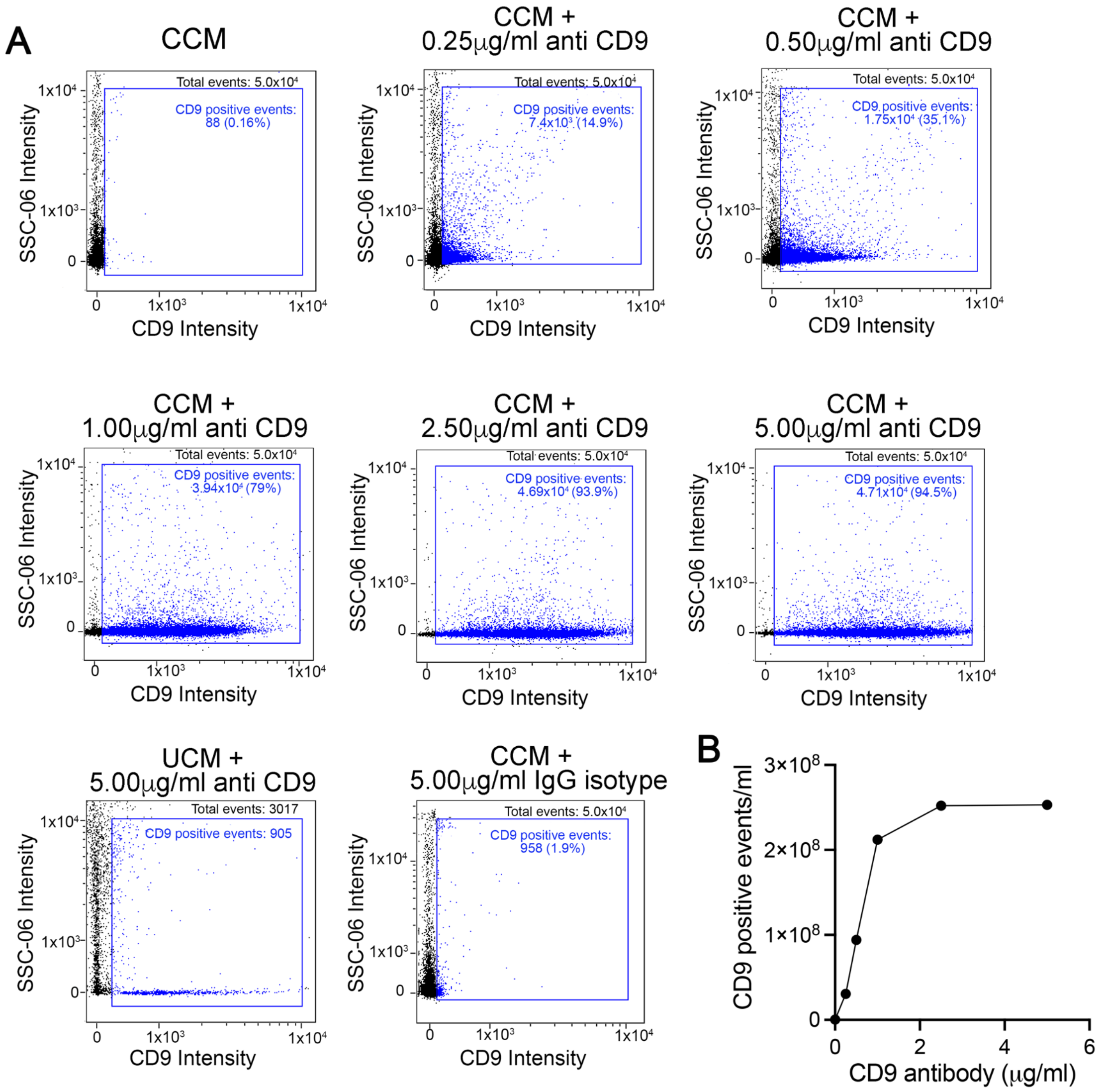


**Supplementary Fig. 1 - Titration of anti CD9 antibody:** The cleared conditioned medium was diluted 1:1 with PBS. Different volumes of a dilution 1:10 of anti-CD9 antibodies in PBS ranging from 0.5 to 10 µl (corresponding to final concentrations of 0.2 to 5 µg/ml) were added to 100 µl of the diluted samples and incubated in the dark at r.t. for 15 min. The following controls were run in parallel: cleared conditioned medium (CCM) without anti-CD9 antibodies, unconditioned medium (UCM) with 5 µg/ml anti-CD9 and CCM with 5 µg/ml IgG isotype. All volumes were equalized with PBS. A) Representative population dot plots at each antibody concentration. The number of CD9-positive events detected in the UCM with the highest concentration of antibody represents ~1.9% of the CD9-positive events detected in the CCM and it is the same number detected by the IgG isotype. B) Quantification of CD9-positive events. Based on this analysis we use the anti-CD9 antibody at a final concentration of 2.5 µg/ml.

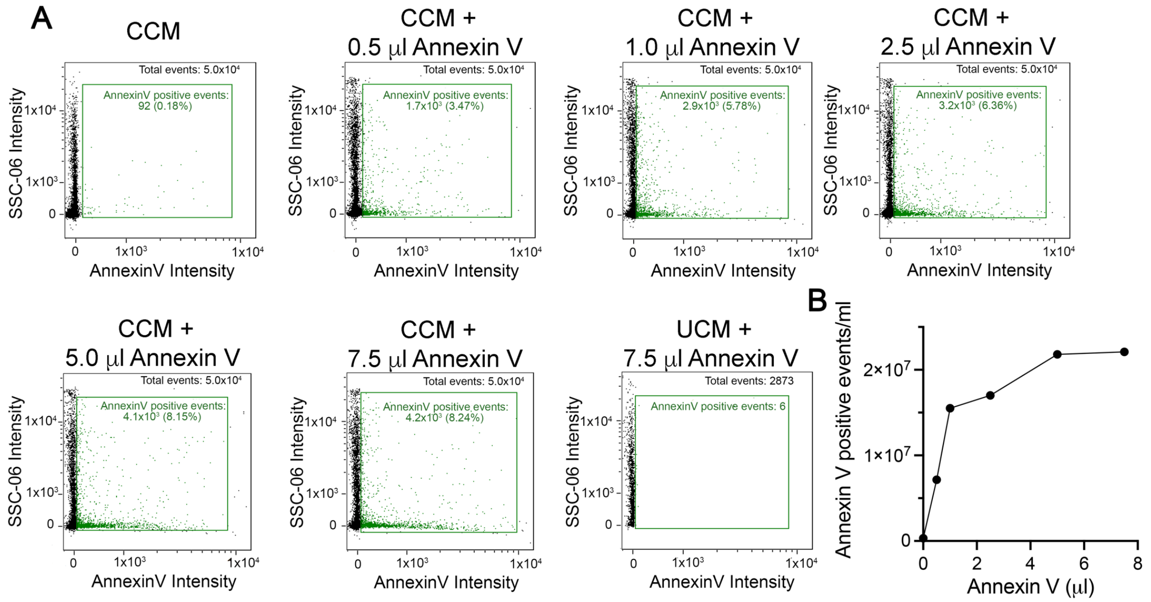


**Supplementary Fig. 2 - Titration of Annexin V:** The cleared conditioned medium was diluted 1:1 with 2X Annexin V binding buffer. Different volumes of Pacific Blue™ Annexin V (concentration proprietary) corresponding to 1:200, 1:100, 1:40, 1:20 and 1:13 dilutions respectively were added to 100 µl of the diluted samples. Controls including CCM without Annexin V, and UCM with 7.5 µl Annexin V were processed in parallel and compared to labelled EV samples. All volumes were equalized with PBS. A) Representative population dot plots at Annexin V dilution. No Annexin V positive particles were detected in the UCM with the highest concentration of Annexin V. B) Quantification of Annexin V-positive events. Based on this analysis we used Annexin V at a final dilution 1/20.

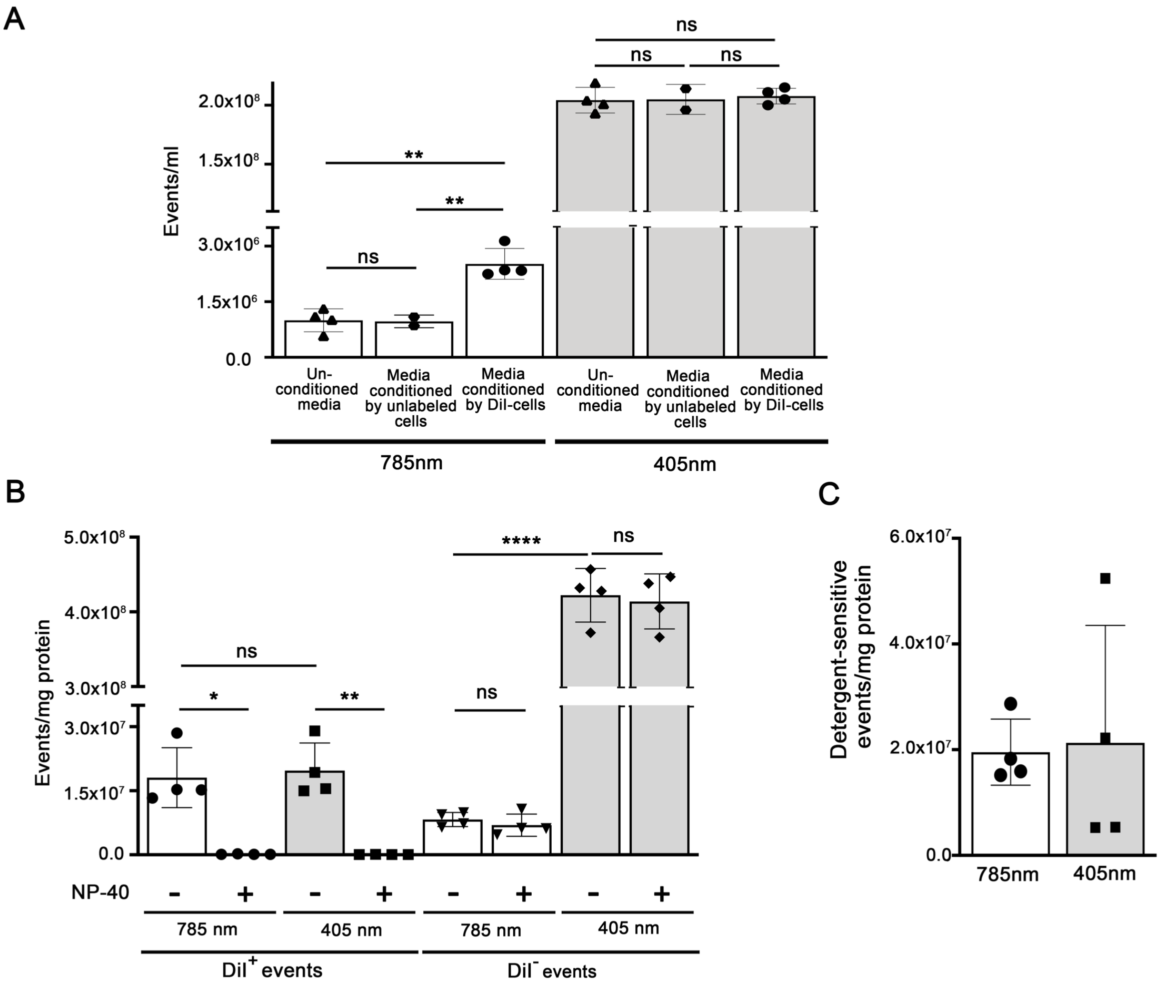


**Supplementary Fig.3 - IFC detection of EVs at different SSC detection laser wavelengths.** **(A)** Quantification of the total number of particles (unlabeled and DiI-labeled) in control unconditioned medium and in the cleared conditioned medium from unlabeled or DiI-labeled cells. Aliquots from the same EV samples were analyzed using a 785 nm or a 405 nm wavelength scatter laser. Significantly more particles are detected with the 405 nm laser, which, however, detects a similar number of particles in the conditioned and the unconditioned control media. Values are means ± SD of 2-4 experiments. One-way ANOVA followed by Sidak's multiple comparisons test separately for each laser group was performed using GraphPad Prism version 9.1.2 for Mac, GraphPad Software, San Diego, California USA, www.graphpad.com. ***p*< 0.01, ns= not significant. **(B)** Both the 785nm and the 405nm scatter lasers allow to detect a similar number of DiI-positive membrane-bound particles sensitive to lysis with NP-40. The majority of particles detected by the 405 nm laser are DiI-negative and not sensitive to lysis with NP-40, suggesting that they are not EVs. Data are mean values ± SD of 4 independent experiments. Paired t-test was applied to assess the statistical significance of the difference between untreated and NP40-treated samples using GraphPad Prism version 9.1.2 for Mac, GraphPad Software, San Diego, California USA, www.graphpad.com". ****p<0.0001, ***p<0.001, **p<0.01 (Paired *t*-test. ***p*< 0.01, ****p*< 0.001, ns= not significant

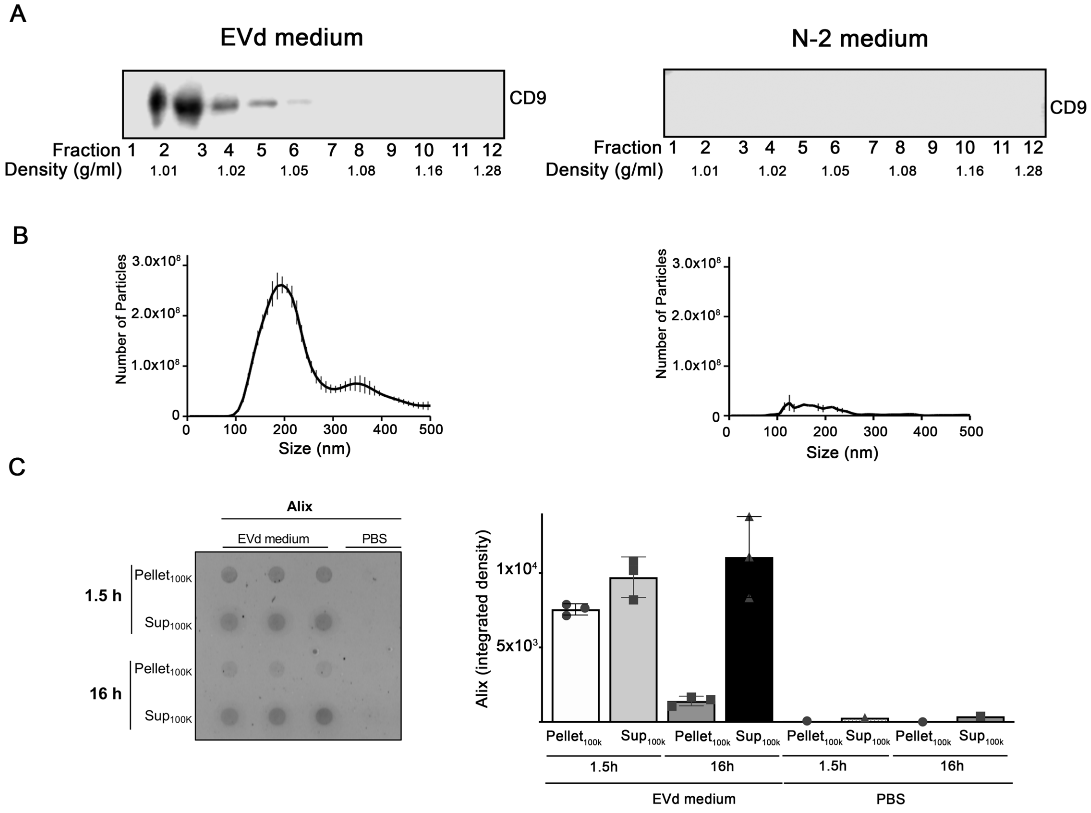


**Supplementary Fig. 4 – EV detection in unconditioned media.** Unconditioned OptiMEM:DMEM (1:1) medium supplemented with either EV-depleted serum (EVd) or N-2 serum-free supplement (N-2) were fractionated by ultracentrifugation in iodixanol gradient and analyzed for the presence of contaminating EV particles that could affect data interpretation in experiments where the medium is conditioned by cells. **(A)** Immunoblotting of fractions probed with an antibody against the EV marker CD9. CD9 was detected in light fractions from EVd medium, indicating the presence of EVs, but not in N-2 medium. **(B)** NTA analysis confirmed the presence of EV particles in the EVd medium. **(C)** EVd medium and control PBS were subjected to UC for 1.5h. The supernatant was centrifuged again at 100,000 x g for 16h. The dot blot with anti-Alix antibodies shows the presence of Alix in both pellet and supernatant from both round of centrifugation, indicating that the majority of EVs in the EVd remain in suspension even after 16h of ultracentrifugation. The graph on the right is the quantification of the signal for Alix in triplicates from one experiment.

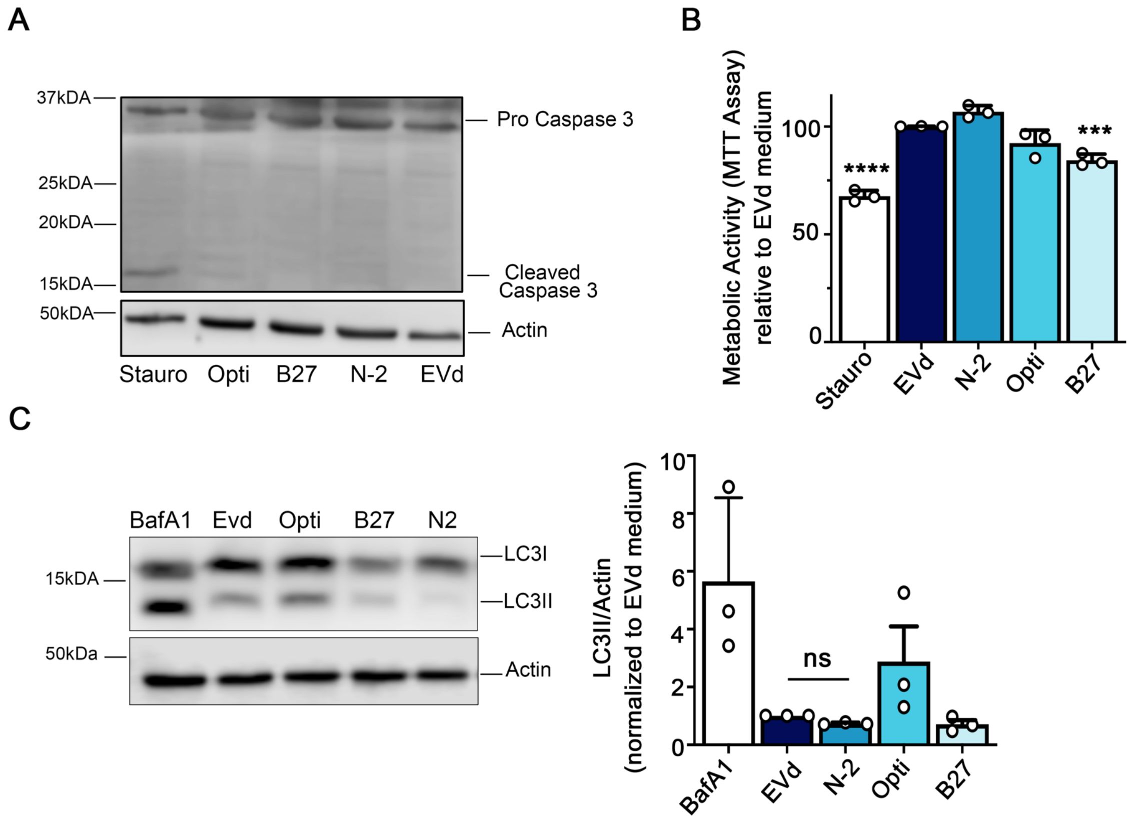


**Supplementary Fig.5: Effects of different culture media on cell viability, metabolism, and autophagy.** N2a cells were cultured in the following media for 24 hours: OptiMEM:DMEM (1:1) supplemented with 10% EV-depleted serum (EVd), serum-free OptiMEM (Opti), serum-free OptiMEM:DMEM (1:1) supplemented with B27 (B27), serum-free OptiMEM:DMEM (1:1) supplemented with N-2 (N-2). **(A)** Caspase 3 activation was measured by immunoblotting to assess whether different media affect cell viability. Cells treated with 10µM staurosporine (Stauro) for 4 h were used as a positive control for activation of caspase 3. **(B)** Cell metabolic activity was determined with the MTT assay. Data were normalized to the metabolic activity of cells cultured in EVd medium. Bars are mean values ± SD of 3 experiments performed in triplicates. One-way ANOVA followed by Dunnett's multiple comparisons test was performed using GraphPad Prism version 9.1.2 for Mac, GraphPad Software, San Diego, California USA, www.graphpad.com ***p<0.001, ****p<0.0001. (**C, D)** Autophagy was assessed by immunoblotting with anti-LC3II antibodies. Cells treated with 150nM bafilomycin (BafA1) were used as a positive control for LC3II accumulation. Data were normalized over cells cultured in EVd medium. Bars represent mean values ± SD of 3 experiments.

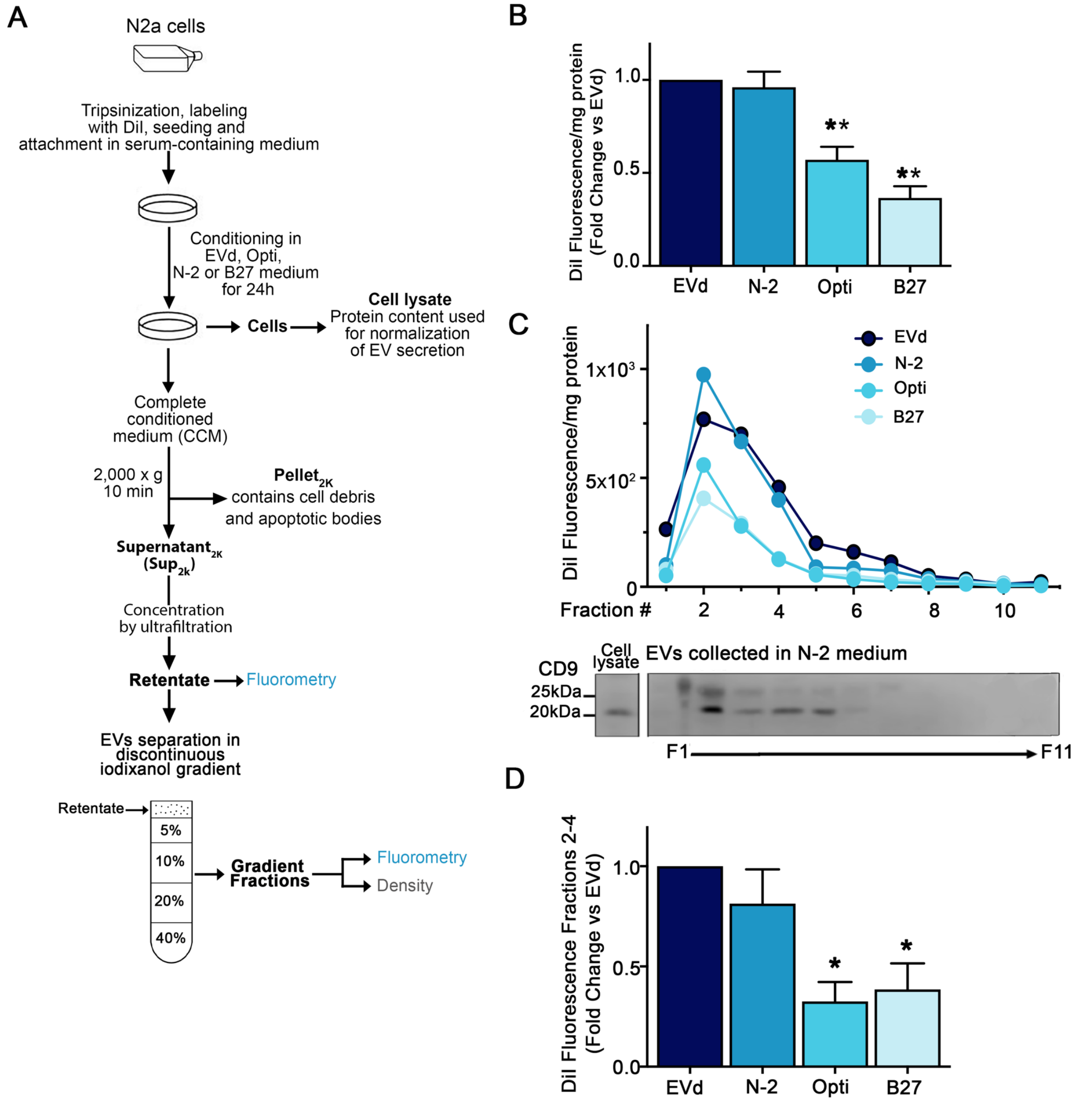


**Supplementary Fig. 6. Effect of culture media on EV secretion**. (**A**) Schematic of EV separation by iodixanol gradient centrifugation. N2a cells were labeled with DiI and plated in complete medium. After 24h, EVs were collected in different media (EVd, N-2, Opti, B27) for 24 h and fractionated on iodixanol gradient. **(B).** DiI fluorescence (EVs) was measured in the concentrated cleared conditioned medium (retentate) and normalized to protein content in the corresponding cell lysate. Results were expressed as fold change vs DiI fluorescence of EVs collected in EVd. The bar graph shows the mean fluorescence values ± SD of 3 experiments. **(C)** After centrifugation in iodixanol density gradient, DiI fluorescence was measured in each fraction and values were normalized to the protein content of the corresponding cell lysate. In parallel, fraction density was determined. One representative experiment out of 3 is shown. The density of each fraction is indicated at the bottom. The immunoblot shows that the majority of EVs is in fractions 2-4, based on the distribution of the EV marker CD9. (**D**). Total EV fluorescence in fractions 2-4 was determined by calculating the area under the curve. Results were expressed as mean fold change values over EVd samples ± SD. N=3. One-way ANOVA followed by Dunnett's multiple comparisons test was performed using GraphPad Prism version 9.1.2 for Mac, GraphPad Software, San Diego, California USA, www.graphpad.com. $^{*}p<0.05$, $^{**}p<0.01$.

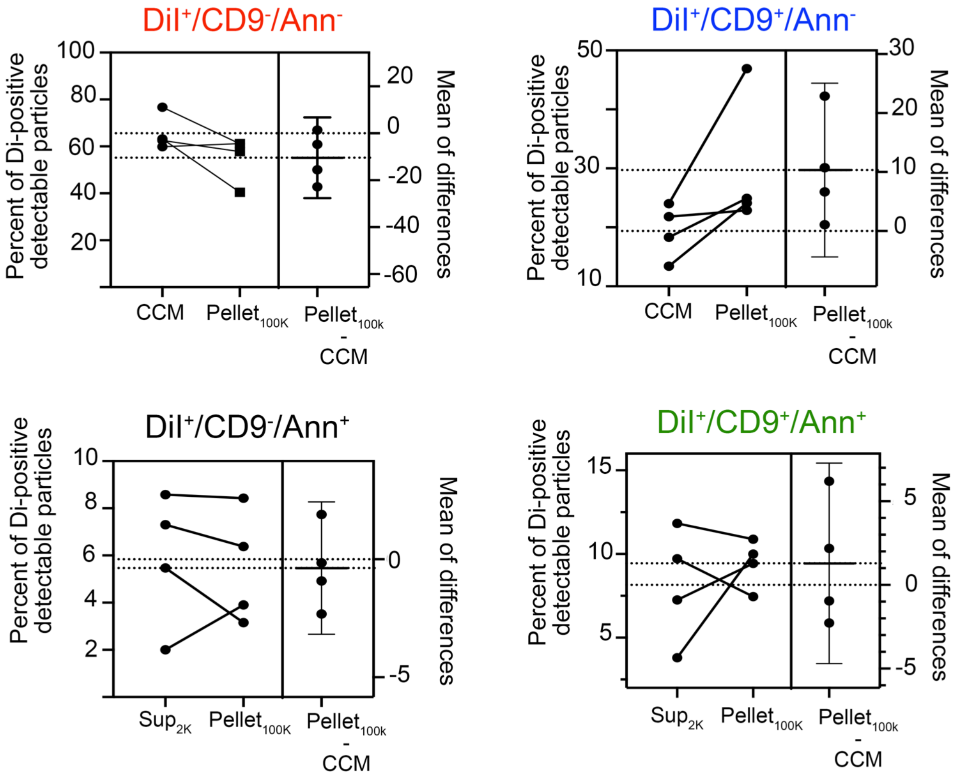


**Supplementary Fig. 7: Comparison of different EVs populations present in the cleared conditioned medium (CCM) and in the Pellet$_{100K}$.** EVs were collected from DiI-labeled N2a and analyzed in the cleared conditioned medium or after isolation by UC. The clear conditioned medium preparations were adjusted based on protein content of donor cells before separation by UC. Clear conditioned medium and Pellet$_{100K}$ preparations were labeled *in vitro* using anti-CD9 antibodies and Annexin V. Quantification of the relative abundance of EV subpopulations in the cleared conditioned medium and in the Pellet$_{100K}$ was performed by IFC. Data are mean percent values of detectable vesicles ± SD of 4 independent experiments. Paired t-test and estimation plot analysis were performed using GraphPad Prism version 9.1.2 for Mac, GraphPad Software, San Diego, California USA, www.graphpad.com. The left panel in each plot shows data for individual experiments as percent values of DiI-positive detectable vesicles. The right panel shows the mean difference between the two groups and the 95% confidence interval of this mean.
No significant differences between groups were found.

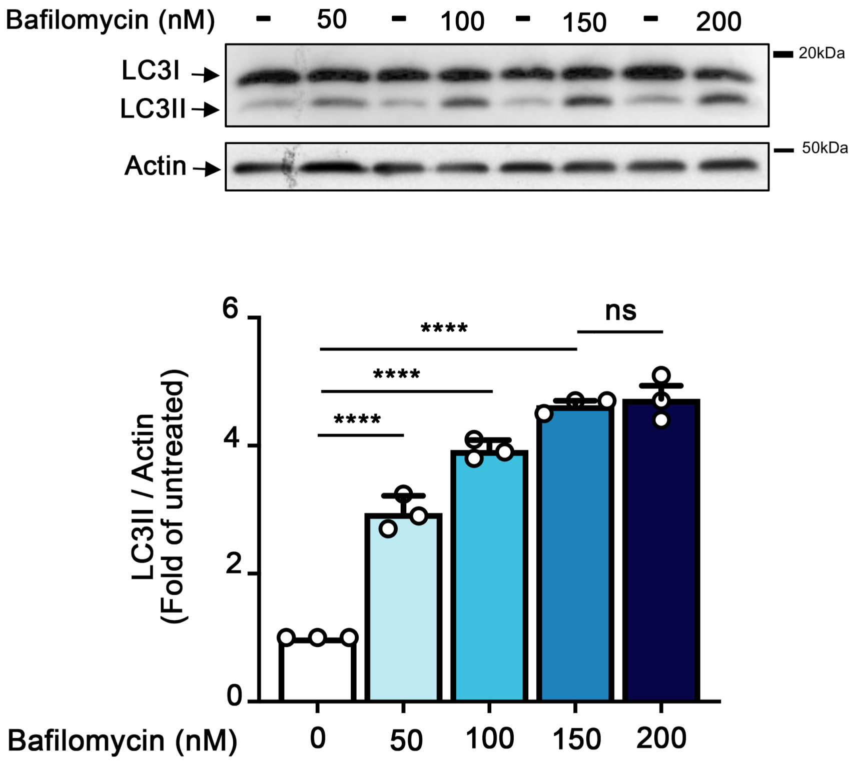


**Supplementary Fig. 8: Blockade of autophagy by Bafilomycin.** N2a cells were treated with vehicle or Baf at different concentrations for 4 h to determine the conditions required to achieve maximum blockade of autophagy. Baf prevents lysosomal acidification and inhibits fusion of autophagosomes/amphisomes with lysosomes. Thus Baf blocks LC3-II degradation. Data were normalized over untreated cells. Bars represent mean values ± SD of 3 experiments.. One-way ANOVA followed by Dunnett's multiple comparisons test was performed using GraphPad Prism version 9.1.2 for Mac, GraphPad Software, San Diego, California USA, www.graphpad.com. ****$p$<0.0001

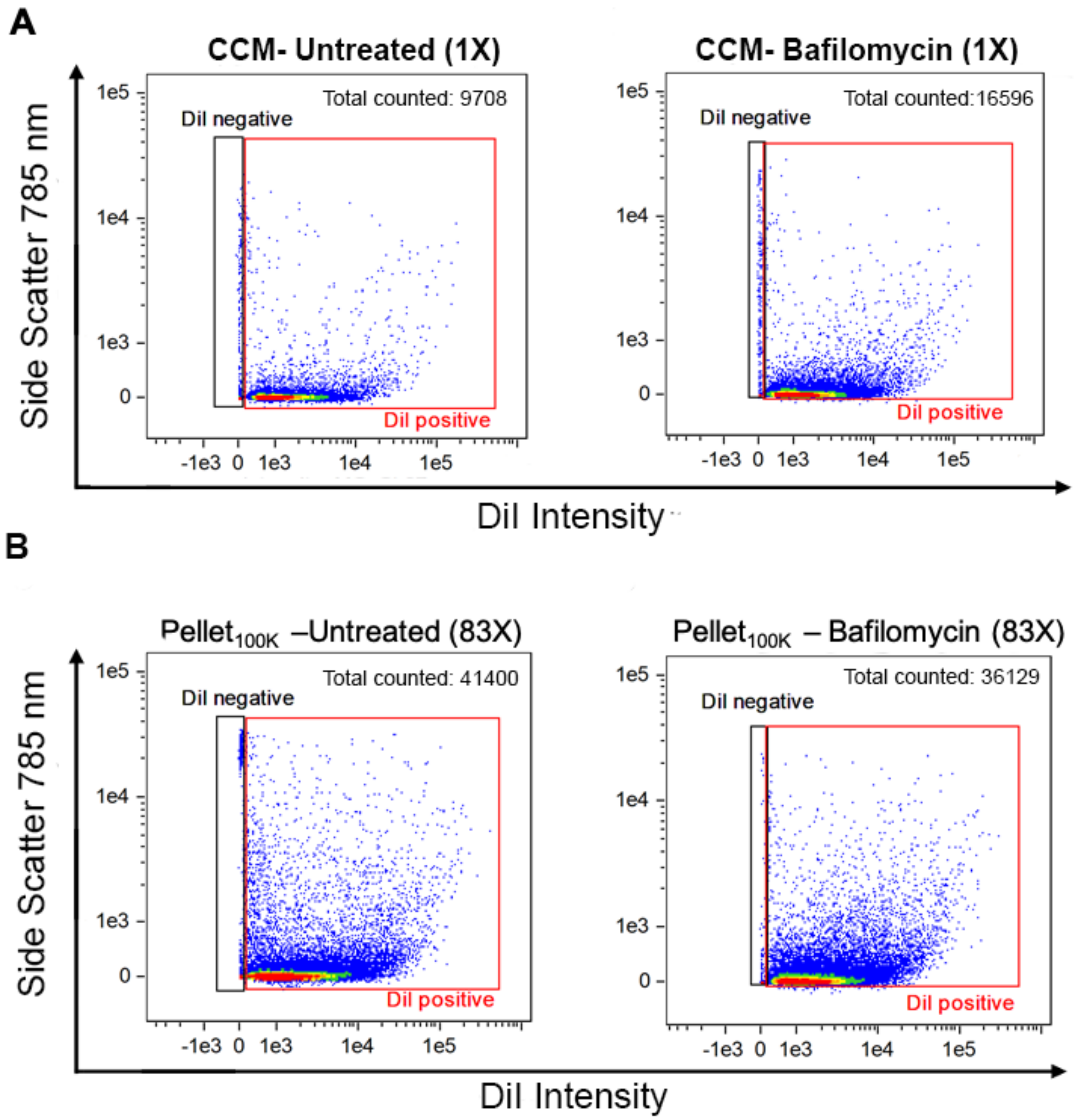


**Supplementary Fig. 9 - Effect of bafilomycin on EV release measured in the cleared conditioned medium and in the Pellet$_{100K}$.** EVs were collected from DiI-labeled N2a cells treated with Baf (150 nM) or vehicle for 4 h and analyzed in the cleared conditioned medium or after isolation by UC. Clear conditioned medium preparations were adjusted based on protein content of donor cells before isolation by UC and analysis by IFC. Representative IFC dot plots are shown. Numbers in parenthesis indicate the concentration factor in each sample. All samples were counted for the same time (10 min).

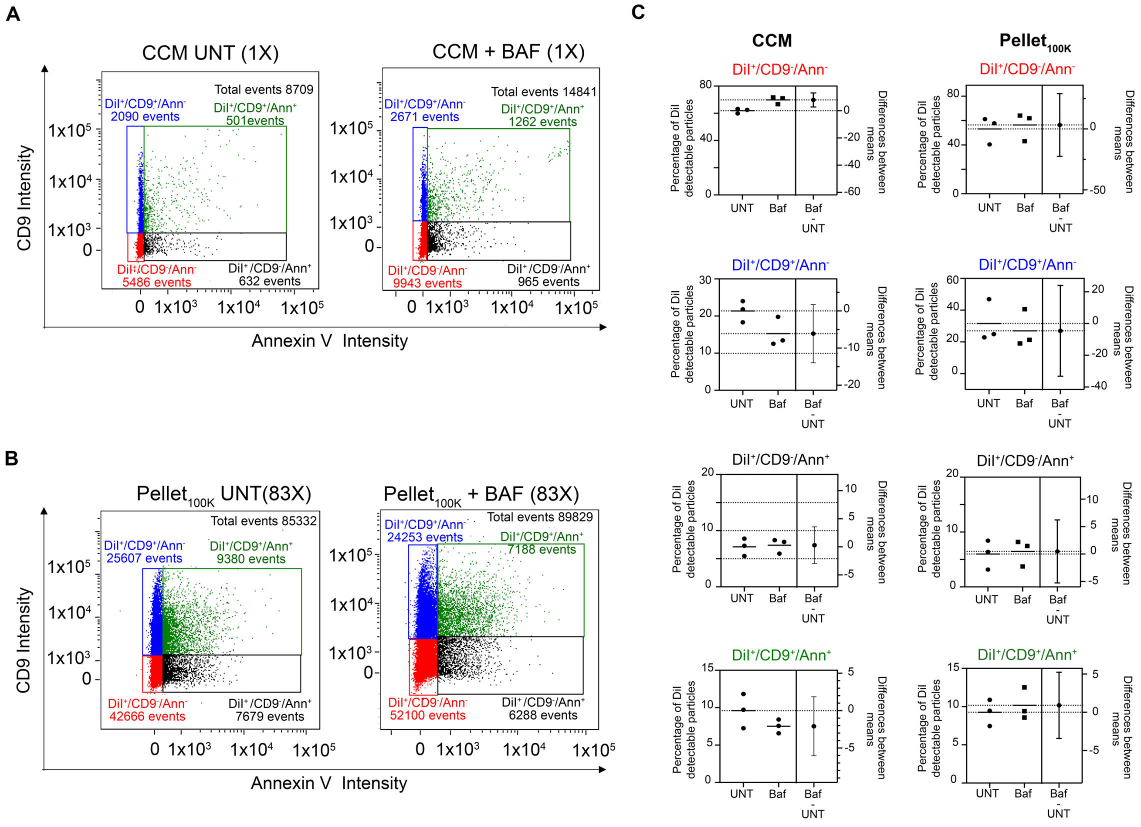


**Supplementary Fig. 10: Effect of bafilomycin on release of different EV populations measured in the cleared conditioned medium and in the Pellet$_{100}$.** EVs were collected from DiI-labeled N2a cells treated with Baf (150 nM) or vehicle for 4 h and analyzed in the cleared conditioned medium or after isolation by UC. Clear conditioned medium preparations were adjusted based on protein content of donor cells before separation by UC. Clear conditioned medium (CCM)and Pellet$_{100K}$ preparations were labeled *in vitro* using anti-CD9 antibodies and Annexin V. Quantification of the relative abundance of EV subpopulations was performed by IFC. **A**) Dotplots of the effect of Baf tested in CCM. **B**) Dotplots of the effect of Baf tested in the Pellet$_{100K}$. **C**) Quantification of the effect of bafilomycin on the total number of each EV population in the CCM (left plots) and in the Pellet$_{100K}$ (right plots). Data are mean percent values ±SD for three independent experiments. Significance determined by unpaired t-test and estimation plot analysis were performed using GraphPad Prism version 9.1.2 for Mac, GraphPad Software, San Diego, California USA, www.graphpad.com. *p<0.05. The left panel in each plot shows data for individual experiments as percent values of detectable vesicles and their mean. The right panel shows the effect size (difference between means) and its 95% confidence interval. The confidence interval for the population DiI$^{+}$/CD9$^{-}$/Ann$^{-}$ in the CCM excludes 0, indicating that the difference is significant.
